# A corticostriatal circuit updates subjective beliefs about latent task states

**DOI:** 10.64898/2026.03.12.711369

**Authors:** Margaret L. DeMaegd, David Hocker, Harsha Gurnani, Mitzi Adler-Wachter, Jonny Schindler, Shannon S. Schiereck, Cristina Savin, Christine M. Constantinople

## Abstract

Beliefs about states of the world profoundly impact decision-making and learning, but little is known about how neural circuits represent and update beliefs. We performed projection-specific recordings and perturbations from neurons in the orbitofrontal cortex (OFC) projecting to the intermediate or rostral caudate putamen (CPi/CPr) in rats performing a task with hidden reward states. Stimulating OFC→CPi neurons biased rats’ beliefs towards high reward states. Recordings from optogenetically-tagged OFC→CPi neurons showed that they encoded categorical evidence for high reward states, shaped by a saturating non-linearity in neural responses. Downstream neurons could, in principle, decode the full belief distribution over reward states as rats deliberated about decisions. Finally, projection-specific perturbations disrupted encoding of hidden states within OFC. These findings reveal the circuit implementation of a core cognitive computation, updating subjective beliefs about abstract latent states of the environment.

## Main Text

Inference is a core computation that is important for many cognitive functions, including perception, motor control, and decision-making (*1*). In partially observable environments, state inference is required for forward mental simulations, planning, or goal-directed decision-making (*2–4*). Uncertainty about the current state can powerfully impact behavior, including by dynamically adjusting the rate at which animals learn from new experiences (*5–9*). Moreover, aberrant belief updating is a hallmark of schizophrenia and other neuropsychiatric disorders (*10–12*). While the orbitofrontal cortex (OFC) has been implicated in inferring values or partially observable task contingencies (*13–18*), how the connectivity patterns and dynamics of synaptic inputs in downstream circuits mediate computations for inference is unclear. Here, we uncover the circuit mechanisms by which OFC neurons that target the intermediate caudate putamen (CPi) represent and causally update beliefs about latent task states in rats performing a rich decision-making task.

## Results

### OFC neurons target CPi and CPr

We trained rats to perform a temporal wagering task in which they revealed how much they valued different amounts of water rewards based on how long they were willing to wait for them (*8*) (Fig. 1A). When rats initiated trials, the frequency of an auditory cue indicated the volume of water reward offered on that trial (5, 10, 20, 40, or 80 µL). At the offset of the cue, the reward was assigned randomly to one of the two side ports, indicated by an LED. The rat could wait for an uncued and unpredictable delay to obtain the reward, or could opt out by poking in the other side port at any time to start a new trial. On 15-25% of trials, rewards were withheld to assess how long rats were willing to wait for them (catch trials), providing a behavioral measure of their subjective value of the reward.

**Figure 1:**
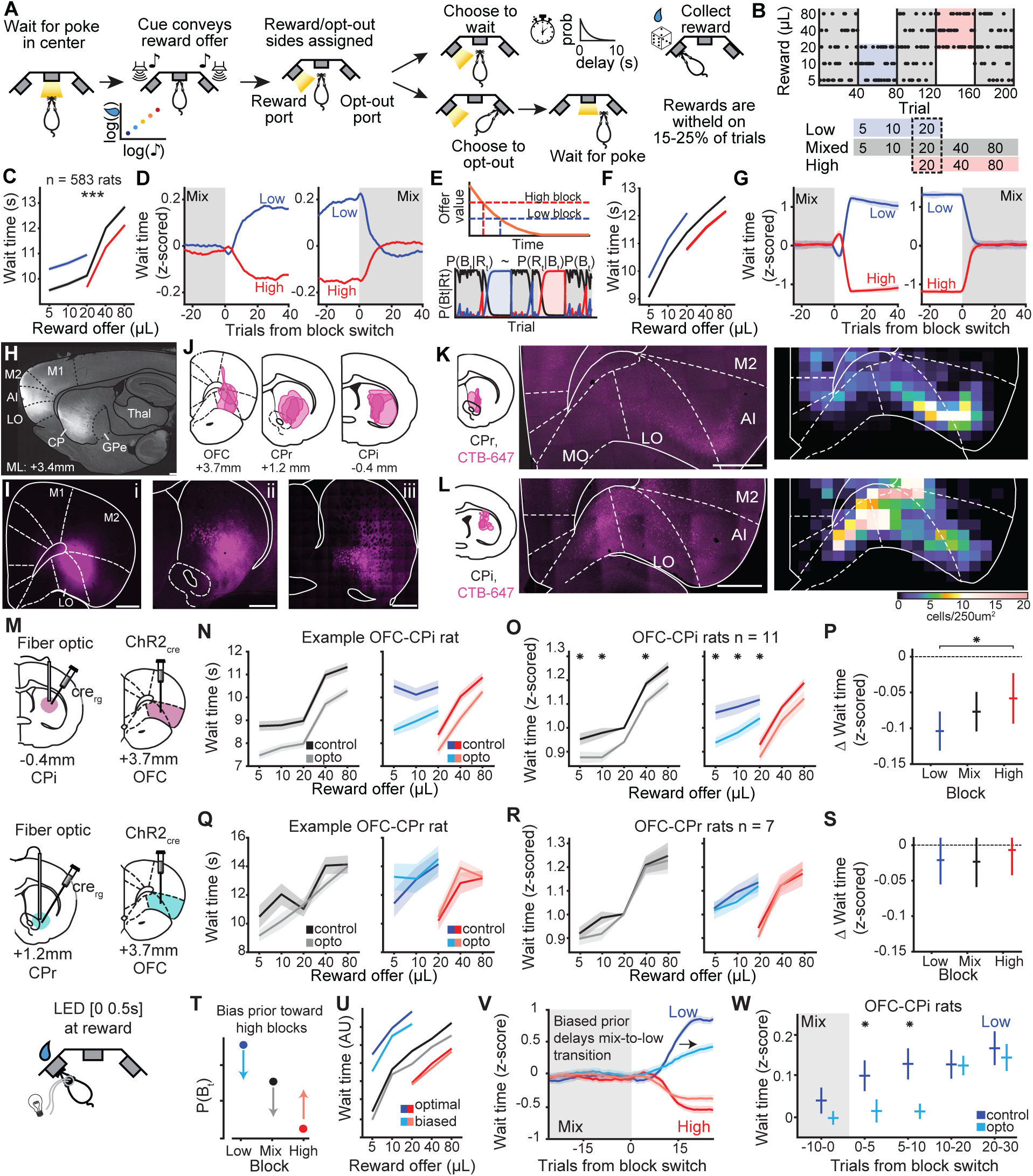
OFC→CPi neurons bias beliefs about hidden states. **A.** Schematic of behavioural paradigm. **B.** Block structure of the task **C.** Mean (+/- sem) wait time on catch trials by reward in each block across all rats. Wilcoxon signed-rank test, *p <<* 0.001. **D.** Mean wait times at block transitions away from (left) or into (right) mixed blocks. **E.** Schematic of the behavioral model, which uses inferential agent that compares a time-varying value function to a block-specific threshold to determine when to opt out. **F.** Mean wait times from the behavioral model. **G.** Model wait times around block transitions. **H.** Sagittal section of a rat brain showing anterograde fluorescent labeling of OFC projections throughout the striatum. Abbrev. M1 - primary motor cortex, M2 - secondary motor cortex, AI - agranular insula, LO - lateral orbitofrontal cortex, CP - caudate putamen, GPe - globus palladas externus, Thal - thalamus. **I.** Coronal section of an example brain at the **i.** OFC injection site Bregma +3.7 mm, **ii.** Bregma 1.2 mm consistent with CPr, and **iii.** Bregma -0.2 mm consistent with CPi. Scale bar = 1 mm. **J.** Overlay of fluorescently labeled regions in N=3 rats. **K.** Overlay of injection sites for retrograde fluorescent labeling of OFC→CPr neurons (left), an example coronal section of the frontal cortex showing labeled somata (middle), and quantification of mean somata across all animals N = 4 (right). **L.** Retrograde fluorescent labeling of OFC→CPi neurons as in K. **M.** Schematic of viral and implant sites in targeting OFC→CPi (top) or OFC →CPr neurons (middle), and LED stimulation which occurred for 500ms aligned to reward delivery on 40% of trials (bottom). **N.** Mean (+/- sem) wait times of an example rat on sessions with LED stimulation targeting OFC→CPi neurons (“opto”, lighter colors) and those without (“control”, darker colors). Data is separated into mixed blocks (left), and low and high blocks (right). **O.** Mean z-scored wait time across all rats as in B. Wilcoxon signed rank test, mixed block: 5 µL *p* = 0.0244, 10 µL *p* = 0.0098, 20 µL *p* = 0.0830, 40 µL *p* = 0.0322, 80 µL *p* = 0.0830, low block: 5 µL *p* = 0.0049, 10 µL *p* = 0.0098, 20 µL *p* = 0.0420, high block: 20 µL *p* = 0.3652, 40 µL *p* = 0.2783, 80 µL *p* = 0.1016. **P.** Mean difference between optogenetic stimulation and control session wait times averaged across all volumes for each block, Friedman test indicating within subject differences *p* = 0.0021, Wilcoxon signed-rank post hoc comparison between low and high block optogenetic modulation, low vs. mix *p* = 0.1607, low vs. high *p* = 0.0084, high vs. mix *p* = 0.1266. **Q-S.** Equivalent figures to N-P in which LED stimulation targeted OFC→CPr neurons. Mean z-scored wait times were not significantly different, Wilcoxon signed rank test, mixed block: 5 µL *p* = 0.6875, 10 µL *p* = 0.2969, 20 µL *p* = 0.9375, 40 µL *p* = 0.8125, 80 µL *p* = 0.6875, low block: 5 µL *p* = 0.8125, 10 µL *p* = 0.3750, 20 µL *p* = 0.8125, high block: 20 µL *p* = 0.3750, 40 µL *p* = 0.9375, 80 µL *p* = 1, nor were mean differences between optogenetic stimulation and control session wait times, Wilcoxon signed-rank post hoc comparison between low and high block optogenetic modulation, low vs. mix *p* = 0.7943, low vs. high *p* = 0.7151, high vs. mix *p* = 0.4140. **T.** Schematic of the biased prior applied to a single trial where the model’s belief favors a low block; the prior biases the model towards high block beliefs and away from low and mixed block beliefs. **U.** Model wait times with an optimal or biased prior. **V.** Model wait times around block transitions. Biased prior delays behavioral changes at mixed→low block transitions. **W.** Mean wait time across all animals in control and optogenetic stimulation sessions around mixed→low block transitions grouped into 5 trial bins. Friedman test, *p <* 0.0247, Wilcoxon signed-rank post hoc test, trials [-10 0] *p* = 0.1602, trials [0 5] *p* = 0.0337, trials [5 10] *p* = 0.0161, trials [10 20] *p* = 0.4492, trials [20 30] *p* = 0.4155.

Rats experienced hidden states, or blocks of trials in which they were presented with low (5, 10, or 20 µL) or high (20, 40, or 80 µL) volumes of water rewards, interleaved with mixed blocks which offered all rewards (Fig. 1B). Wait times were sensitive to the hidden reward states (Average responses shown in Fig. 1C,D, and individual examples in fig. S 1). Rats’ behavior is well-accounted for by a model in which they compare the value of waiting to a threshold that reflects the opportunity cost, or average reward in each block (*8*, *18*, *19*) (Fig. 1E). The model uses Bayes’ rule to infer the most likely block, and implements a fixed, block-specific wait time threshold based on that state inference. We note that the shape of the wait time curves somewhat differs between the rats and the model, due to the model’s assumption that subjective value reflects the binary logarithm of the reward volume. We view this as a minor difference, as the focus of the present study is on the modulation of wait times by the hidden reward states. In that respect, the inferential model reproduces myriad aspects of behavior (*8*, *18*) (Fig. 1F,G). We previously found that inference in this task requires the OFC, and inactivating OFC impairs rats’ ability to update their beliefs about the block (*18*). Here, we sought to characterize how OFC projection neurons contribute to the belief updating computation.

Cortical innervation of the striatum, the input structure of the basal ganglia, is extensive and topographic. Mesoscale projection mapping studies in the mouse have divided the caudate putamen into rostral extreme (CPre), rostral (CPr), intermediate (CPi), caudal (CPc), and caudal extreme (CPce) subregions (*20*, *21*). Subregions can be further segmented into “communities” based on cortical and thalamic innervation patterns (*20*, *22*). To characterize OFC innervation of CP subregions in the rat, we injected AAV-CB7-mCherry or AAV-hSyn-mCherry in the lateral OFC (Fig. 1H,I). Consistent with past studies of rat OFC, we targeted LO and the agranular insular cortex (AI) (*18*, *23–28*). We identified terminal fields in the CPr and CPi (Fig. 1J). At the level of the CPr, lateral OFC axons innervated the intermediate ventral community, consistent with ventrolateral OFC projections in the mouse (*20*). At the level of CPi, we observed innervation of the ventrolateral community. This is in contrast with findings in the mouse, which have reported OFC innervation of dorsomedial subregions, although anterograde injections in those studies were more medial, and appeared more anterior than our injections based on anatomical landmarks (*20*, *21*). Species differences between mice and rats could also contribute to this discrepancy.

We next injected the retrograde fluorescent tracers CTB-Alexa 488 or CTB-Alexa 647 in the intermediate ventral CPr or ventrolateral CPi (Fig. 1K,L). Quantification of retrogradely-labeled cell bodies in OFC revealed that the lateral LO and AI were enriched with neurons that projected to the CPr, and this projection class straddled the putative boundary between these regions. In contrast, neurons projecting to CPi tended to be localized in LO and more medial orbital regions.

We next sought to characterize the function of OFC→CPr and OFC→CPi neurons in the temporal wagering task.

### OFC→CPi neurons bias beliefs

We bilaterally injected a retrograde Cre virus in the CPi or CPr and a Cre-inducible excitatory opsin (ChR2.0) in the OFC, and implanted a tapered fiber optic among OFC→CP axon terminals in the striatum (Fig. 1M). In optogenetic sessions, we delivered blue light (465nm) for 500 ms aligned to the time of reward delivery on 40% of trials. This manipulation caused rats to wait less time for rewards in each of the blocks (Fig. 1N,O). This optogenetic effect persisted over trials, as light was delivered at the time of reward, but the behavioral effect was apparent on trials in which rats opted out and received no reward, which are by definition different trials from the stimulation trials. Notably, the optogenetic effect size depended on the block, with the largest effect in low blocks, an intermediate effect in mixed blocks, and the weakest effect in high blocks (Fig. 1P).

The block-dependent effects were supported by a Friedman test indicating within subject differences, and by the relative fraction of reward offers with significant wait time differences in each of the blocks (Fig. 1O). In contrast, optogenetically stimulating OFC→CPr neurons at reward delivery failed to change how long rats were willing to wait for rewards compared to control sessions (Fig. 1Q-S). Comparisons between the change in wait times caused by stimulation of OFC→CPi and OFC→CPr neurons indicate a significantly greater effect for OFC→CPi targeted animals (Wilcoxon rank sum test, p=0.0079). Altogether, these data indicate that these corticostriatal pathways are functionally dissociable, with OFC→CPi having a privileged role over OFC→CPr neurons for modulating wait times according to the block.

We sought to use our behavioral model to generate a hypothesis about the cognitive process that was disrupted by optogenetic stimulation. The model estimates the most likely block (*P* (*B_t_*|*R_t_*)) by combining the likelihood of observing the current reward in that block (*P* (*R_t_*|*B_t_*)) with the prior belief about the block (*P* (*B_t_*)), where the prior is recursively updated over trials. We found that if we implemented a biased prior, such that the model is biased to believe it is in a high block and against believing it is in a mixed or low block, this reduces how long the model is willing to wait, with the largest change in low blocks, intermediate change in mixed blocks, and weakest change in high blocks (Fig. 1T,U). Intuitively, the biased prior induces mistaken inferences in the model, increasing the probability that it will use the opportunity cost or wait time threshold associated with a high block. Because the opportunity cost of the low block is the most different from a high block, the strongest effect on wait times is in low blocks, and intermediate in mixed blocks. Even an ideal Bayesian observer would make mistaken inferences on about 10% of trials (*8*); the biased prior eliminates mistaken inferences in high blocks, thereby also modestly reducing wait times in those blocks.

As an additional test, we simulated the behavior of the model with an optimal versus biased prior, and visualized the changes in wait times at different block transitions. Because the wait times are modulated by both the reward volumes and the blocks, we first z-scored the wait times for each reward independently, before pooling z-scored wait times over different reward volumes, to isolate contextual effects from the effects of reward volume on wait times (*18*). The prior that was biased towards high blocks made the model especially resistant to accepting that it had transitioned from a mixed into a low block, predicting that wait times should be slower to change specifically at this transition but not others (Fig. 1V). To test this prediction, we compared the rats’ z-scored wait times in different bins of five trials at transitions from mixed into low blocks, for control and optogenetic sessions. Consistent with the model, wait times in optogenetic sessions were slower to change following transitions into low blocks (Fig. 1W). Similar effects were not observed at other block transitions, consistent with the model prediction (fig. S 2).

We tested whether other model parameters could reproduce these patterns of behavioral results (fig. S 3). Increasing the opportunity cost associated with each block did decrease model wait times, but failed to reproduce the block-dependent effect sizes or slower behavioral changes at mixed-to-low block transitions. Modifying the model’s time-varying reward expectations (i.e., the reward hazard rate), or introducing a bias in the perception of rewards towards larger offers, did reproduce block-dependent changes in wait times. However, neither reproduced the slower behavioral changes at mixed-to-low block transitions. In contrast, the model with biased beliefs towards high blocks could reproduce all the patterns of behavioral changes produced by optogenetic stimulation.

### OFC→CPi neurons encode high block evidence

We next sought to characterize the task-related dynamics of OFC→CPi and OFC→CPr neurons. We used a viral strategy to express ChR2.0 in these projection neurons, implanted a fiber optic in the CPi or CPr, and a Neuropixels probe in the OFC (Fig. 2A). We recorded neural activity from the probe as rats performed the task, and at the end of the behavioral session, we antidromically stimulated OFC→CP axons, to identify projection-specific neurons on the basis of collision tests (*29*). At a precise delay, orthodromic and antidromic action potentials should ‘collide’ and, due to refractory sodium channels on either side of the collision, cancel (Fig. 2B,C). These neurons showed highly reliable, short latency responses to LED stimulation, and endogenous and elicited spikes produced similar waveforms (fig. S 4). Leveraging our high-throughput behavioral training approach which yielded large pools of trained subjects for experiments, we recorded from 95 OFC neurons projecting to the CPr or CPi, identified from collision tests.

**Figure 2:**
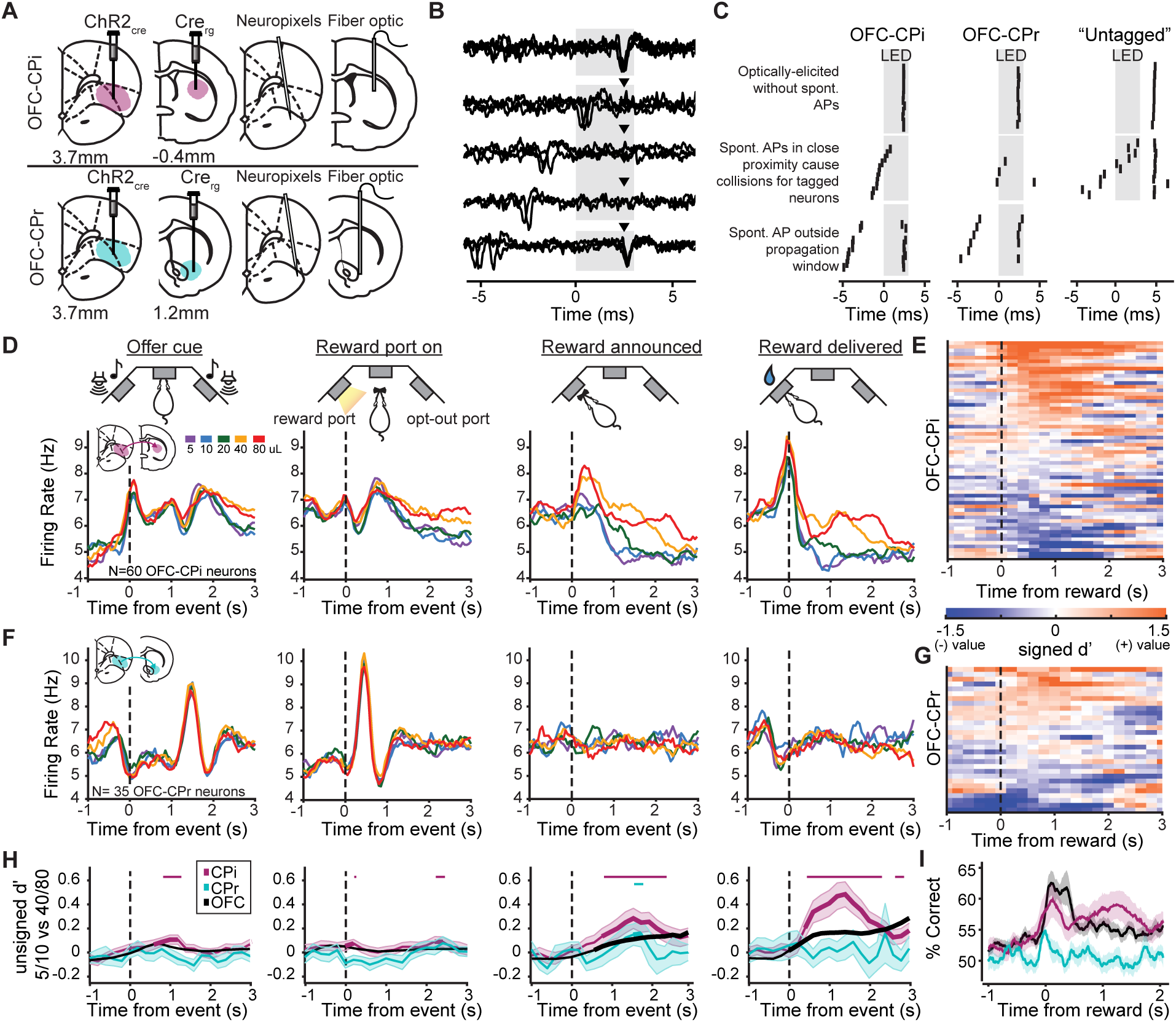
OFC→CPi neurons, but not OFC→CPr neurons, convey high block evidence to striatum. **A.** Schematic of viral and implant strategies to target projection-specific neurons. **B.** Waveform overlays of spikes from an example OFC→CPi neuron aligned to LED onset and sorted by the timing of the last action potential prior to the LED onset. Arrows indicate the expected timing of LED elicited spikes. **C.** Raster plots of spikes from three example cells recorded in the OFC which either pass collision tests and are therefore projection neurons to the CPi and CPr (left and middle respectively) or do not pass the collision test are therefore do not directly project to the striatum and labeled “untagged.” **D.** Mean reward-dependent PSTH of OFC→CPi neurons aligned to each task event. N=60 neurons, n=5 rats. For the number of cells per recording session see Supplemental Data Table S1. **E.** Heatmap of signed discriminability index (d’) for trials offering 40/80 versus 5/10 µL in mixed blocks for each OFC→CPi neuron aligned to reward delivery. **F-G.** Same analyses in D,E for OFC→CPr neurons. **H.** Mean (+/- sem) unsigned discriminability index (d’) for trials offering 40/80 versus 5/10 µL in mixed blocks for OFC→CPi (pink), OFC→CPr (teal), and untagged OFC neurons (black). Bolded traces indicate time points where the unsigned d’ is larger than the 95th percentile of the label-shuffled data for that neuronal population. Time points with significant differences (*p <* 0.05, non-parametric permutation test) between OFC→CPi or OFC→CPr and untagged OFC neurons is depicted in matching colors above each PSTH and actual p values for both tests are listed in Supplemental Data Table S2. **I.** Logistic decoding of reward volume (40/80 *µ*L or 5/10 *µ*L) in mixed blocks.

On average, OFC→CPi neurons appeared to encode evidence in favor of high blocks at the time of reward delivery (Fig. 2D), consistent with our optogenetic manipulation of OFC→CPi neurons in which stimulation biased beliefs towards high blocks. Specifically, they tended to have higher firing rates for 40/80 µL compared to 5/10 µL. To determine whether this was a unique feature of this projection class, we computed the mean unsigned discriminability index (d’) for trials offering 40/80 µL versus 5/10 µL in mixed blocks (Methods). OFC→CPi neurons exhibited significantly higher d’ values than OFC→CPr neurons, or the overall untagged OFC population, particularly at the time of reward (Fig. 2F,H). We trained a logistic regression decoder to decode whether rewards in mixed blocks were large (40/80 µL) or small (5/10 µL) based on the firing rates of OFC→CPi neurons, OFC→CPr neurons, or the total OFC population (Fig. 2I). Decoder performance was higher for OFC→CPi compared to OFC→CPr neurons, and comparable to the entire OFC population, which included many thousands of neurons, some of which presumably projected to the CPi. This shows that OFC→CPi and OFC→CPr neurons convey distinct information to the striatum.

40 and 80 µL are different reward volumes but provide identical evidence for high blocks, and 5 and 10 µL similarly provide identical evidence for low blocks (and against high blocks). The fact that the mean firing rates of OFC→CPi neurons was similar for 40/80 µL and 5/10 µL (Fig. 2D) suggests that these neurons encode categorical evidence for updating beliefs about the blocks, as opposed to graded representations of reward volumes *per se*. Because on average, the OFC→CPi neurons have higher activity for high block evidence (i.e., 40/80 µL), increasing activity in this population would induce a downstream integrator to integrate additional evidence for high blocks, thus biasing the rats’ beliefs towards high blocks.

### Mechanisms of the block evidence computation

What mechanism might convert information about graded reward volumes into categorical evidence for belief updating? Notably, there was heterogeneity in the sign of evidence encoding in the OFC→CPi population (Fig. 2E). Visualizing the signed d’ for large versus small rewards across all projection neurons revealed a bias towards positive d’ values, that is, neurons with higher firing rates for large rewards, but some neurons exhibited negative d’, or higher firing rates for small rewards (n=27 neurons significant positive d’, n=12 neurons significant negative d’, n=21 neurons not significant, Methods). We identified neurons whose mean d’ in the [0 1s] window after reward was in the bottom (negative d’) or top (positive d’) quintiles (full distribution shown in fig. S 5A). There were no significant differences in response variance across quintiles (fig. S 5B,C), indicating that differences in d’ reflected mean firing rate differences across rewards. On average, positive d’ neurons appeared to encode preferred 40/80 µL rewards categorically, but exhibited more graded encoding of non-preferred, smaller rewards [5 10 20 µL] (Fig. 3A, left). Similarly, negative d’ neurons encoded preferred 5/10 µL rewards categorically but exhibited more graded responses for non-preferred larger rewards [20 40 80 µL] (Fig. 3B, left). This held if we quantified discriminability for 40 µL *versus* 80 µL and 5 µL *versus* 10 µL in these populations (Fig. 3A,B right). To control for potential motor correlates due to shorter drinking times for smaller rewards, we restricted this analysis to a window during which rats were drinking for even the smallest volumes, the [0 1 s] window after reward. For both groups, rewards that elicited higher firing rates (preferred rewards) were less discriminable than non-preferred rewards.

**Figure 3:**
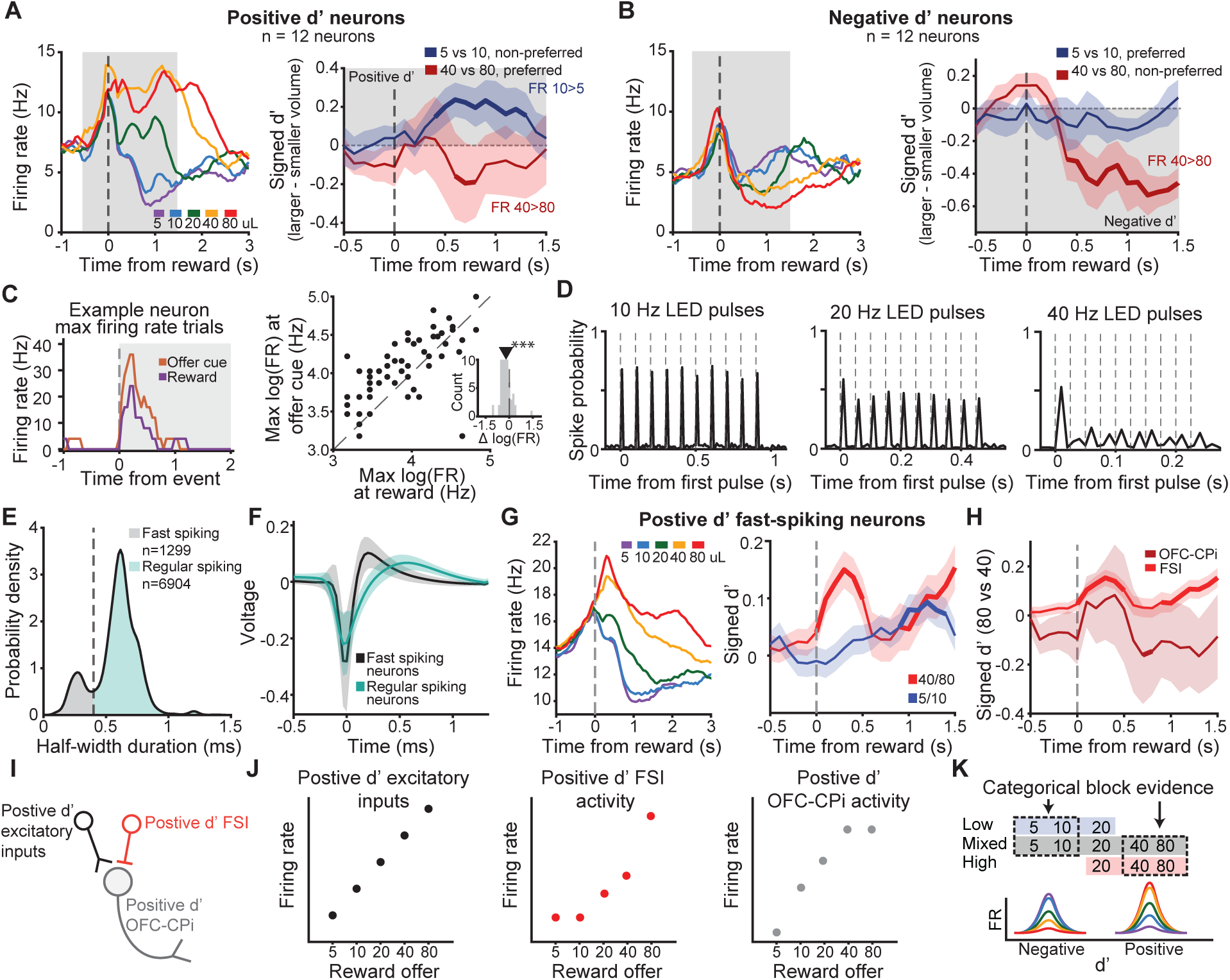
Saturating nonlinearity implements categorical encoding of block evidence in OFC→CPi neurons. **A.** PSTH of positive d’ neurons for different reward volumes and aligned to reward delivery (left). Mean (+/- sem) discriminability between the neurons’ two preferred volumes (reward volumes with the highest firing rate responses; 40 vs. 80 µL) and non-preferred volumes (reward volumes with the lowest firing rate responses; 5 vs. 10 µL). Bolded traces indicate time points where discriminability is larger than the 95th percentile of the label-shuffled data. Grey box indicates d’ values consistent with monotonic reward coding, i.e. higher firing rates for larger volumes. **B.** PSTH of negative d’ neurons for different reward volumes and aligned to reward delivery (left). Mean discriminability of the neurons’ two preferred volumes (5 vs. 10 µL) and non-preferred volumes (40 vs. 80 µL). Bolded traces indicate time points where discrim-inability is larger than the 95th percentile of the label-shuffled data. Grey box indicates d’ values consistent with monotonic reward coding, i.e., higher firing rates for smaller volumes. **C.** PSTH of an example OFC→CPi neuron on the single trials with the highest firing rate during offer cue (orange) and reward delivery (purple). Right: Maximum firing rate at offer cue and reward de-livery for all projection neurons. Inset shows a histogram of the difference in maximum firing rate. **D.** Spike probability aligned to the first LED pulse in a 10 pulse train at 10 Hz (left), 20 Hz (middle), 40 Hz (right) for an example neuron. Dashed lines indicate the onset of each LED pulse in the train. **E.** Distribution of waveform half-width durations. Regular (teal) and putative fast (black) spiking neurons using a half-width spike duration threshold of 0.4 ms. **F.** Mean +/-STD waveform of fast (black) and regular (teal) spiking neurons. **G.** Left: PSTH of positive d’ FSIs sorted by reward volume and aligned to reward delivery. Right: FSI discriminability in encoding 5µL versus 10µL and 40µL versus 80µL rewards. **H** Mean (+/- sem) discriminability for 40µL versus 80µL for positive d’ OFC-CPi and FSI neurons, overlaid. Bolded traces indicate time points where d’ is larger than the 95th percentile of the label-shuffled data. FSIs show higher firing rates for 80µL when OFC→CPi neurons categorically encode rewards. **I.** Schematic of hypothesized mechanism by which linear positive d’ excitatory input and local inhibition from positive d’ FSI target a postive d’ OFC→CPi projection neurons to support categorical evidence of a high block. **J.** Hypothesis: OFC→CPi neurons receive graded excitatory input proportional to reward (left). Local inhibition from fast-spiking interneurons is strongest for large reward volumes (middle) which saturates the peak firing of OFC→CPi neurons in response to larger volumes (right). **K.** Schematic of hypothesized mechanism of block evidence computation. Saturating non-linear reward encoding for the two highest and lowest reward volumes which provide equal evidence in favor of a high or low block respectively. Heterogeneity in the sign of encoding supports categorical encoding of both large and small rewards.

We hypothesized that this effect might reflect a saturating non-linearity in neural responses. We measured the maximum firing rate of each neuron at other task events, including the offer cue and when the side light turned on indicating the reward port. We found that most neurons had higher firing rates at other task epochs, compared to the reward epoch (Fig. 3C,D), indicating that the saturating firing rate for preferred rewards is not a cell-intrinsic maximum, but may involve recurrent circuit dynamics including local inhibition. Indeed, while OFC→CPi neurons were faithfully driven by 10 Hz optogenetic stimulation, they showed reduced ability to follow 20 Hz and 40 Hz pulse trains (Fig. 3C,D), consistent with recruitment of inhibition at higher frequencies (*30–32*).

We next tested whether the responses of putative fast-spiking interneurons (FSIs), identified by their narrow spike widths, were consistent with implementing a saturating nonlinearity (Fig. 3E,F). Putative FSIs showed heterogeneity in reward encoding (fig. S 6), so we again separated FSIs whose mean d’ values after reward were in the top (positive d’) or bottom (negative d’) quintiles. Remarkably, the positive d’ FSIs exhibited the reverse pattern of responses observed in the positive d’ OFC→CPi neurons: responses for preferred rewards were *more discriminable* than for non-preferred rewards (Fig. 3G,H). In principle, stronger inhibition for 80 µL could more strongly suppress positive d’ OFC→CPi neurons on those trials, and if appropriately tuned, result in categorical encoding of 40/80 (Fig. 3I,J). This result did not hold for negative d’ FSIs (fig. S 6). Additional mechanisms, such as saturation of excitatory inputs, synaptic depression, or the involvement of other types of interneurons, may contribute to the block evidence computation, or differentially contribute for positive versus negative d’ OFC→CPi neurons. The results from the positive d’ FSIs provide some of the first evidence for well-defined inhibitory neuron computations in frontal cortex in the context of a rich decision-making task.

Taken together, these findings show that the hallmark of the block evidence computation is that graded representations of reward volumes, which are apparent for non-preferred rewards that elicit lower firing rates in OFC→CPi neurons’ linear dynamic range, are transformed into categorical representations of evidence for belief updating. Heterogeneity in the sign of evidence encoding (i.e., the fact that some neurons have positive d’ values and others have negative d’ values) enables categorical encoding of both large and small rewards at the population level (Fig. 3K).

### Decodable belief distributions

Based on the responses of the OFC→CPi neurons, and our optogenetic results, belief updating appears to occur at the time of reward. We next wondered if the OFC→CPi neurons might represent subjective beliefs about the blocks, perhaps at other times in the trial. The posterior distribution over blocks is a discrete probability distribution with three values on each trial, one for the posterior probability of each block. Since the probabilities must sum to one, being able to decode any two of those probabilities allows one to estimate the third, i.e., know the full probability distribution (Fig. 4A).

**Figure 4:**
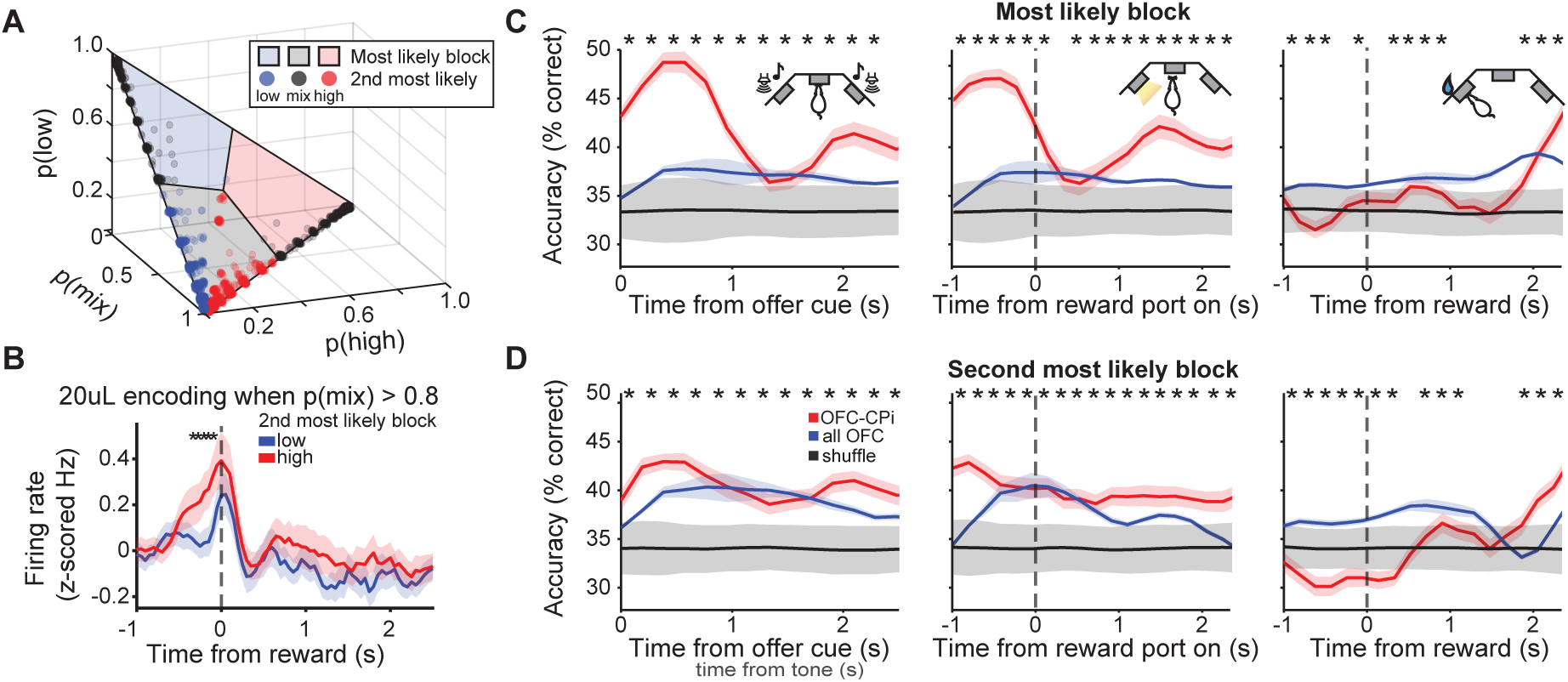
OFC→CPi neurons convey the full belief distribution in other task epochs. **A.** Distribution of most likely and second most likely block from OFC→CPi recording sessions. Block probabilities for each rewarded trial were estimated from the Bayesian model of the task (*8*), are plotted in 3D probability space. The colored regions of the simplex correspond to the most likely block for that trial, and dot colors convey second most likely block. Red = high block, black = mixed block, blue = low block. **B.** Mean (+/- sem) PSTH of OFC→CPi activity conditioned on 20 µL trials in a mixed block in which the behavioral model indicated that the probability of a mixed block was high (P(mix) *>* 0.8) and the second most likely block was high (red) or low (blue) with a low probability (P*<* 0.05). Significant firing rate differences are denoted above the traces (non-parametric permutation test, *p <* 0.05, actual p values listed in Supplemental Data Table S5). **C-D.** Decoding accuracy of OFC→CPi neurons to classify (C) most likely block and (D) second most likely block. Each column denotes decoding at a different epoch within the task. Colored lines represent median decoding over different populations (red = OFC→CPi, blue = all OFC, black = shuffle distribution built by shuffling trials labels of OFC→CPi data.) Error bars for red and blue curves are sem, and error bars for the shuffle distribution are 1.96 standard deviations calculated over 20 shuffles. Significant differences in decoding accuracy between OFC→CPi neurons and a sample-number matched untagged OFC population via rank sum test (*p <* 0.05) is denoted above each timepoint. Data have been smoothed with a causal filter of 3 data points (600ms) for visualization only.

Because the block probabilities are highly correlated with the reward offers in this task, we computed the average firing rate of OFC→CPi neurons for trials offering 20 µL, with similar maximum a posteriori estimates (i.e., probability for the most likely block) but different full posterior distributions. Specifically, we selected trials where the most likely block was a mixed block, with a high probability (*>*= 0.8), and the second most likely block was either a high or a low block with a low probability (*<*0.05). Given that the reward (20 µL) and most likely block (mixed) are all identical on these trials, any differences in firing rates would be consistent with reflecting the full posterior distribution. Trials meeting these strict criteria were rare, and are represented by a subset of red and blue dots in the mixed block sector (black shaded region) in Figure 4A. Despite the limited trials, there was a significant difference in the average firing rates of OFC→CPi neurons across these conditions, prior to reward delivery while the animals waited for rewards (Fig. 4B). This suggests that OFC→CPi neurons represent the full belief distribution as rats are deciding how long to wait.

We additionally used a decoding approach to test whether downstream neurons in the striatum would be able to decode the first and second most likely block, predicted from the Bayesian model, from this projection class. Most optogenetically-tagged neurons were not recorded simultaneously, so we constructed pseudo-populations for a logistic regression decoder, using all rewarded trials, and evaluated performance using leave-one-out cross validation. Because the block identity is highly correlated with reward volumes, we trained and tested the decoder after subtracting out the mean PSTH for each reward offer. It was possible to decode the first and second most likely block from this projection class (using separate decoders) early in the trial, when rats heard the offer cue and as rats waited for rewards, indicating that in principle the CPi could estimate the full posterior distribution over blocks during these epochs (Fig. 4C,D). These results were robust and held if we did not subtract the mean PSTH for each reward offer (fig. S 7). In contrast, while it was possible to decode the first and second most likely block from OFC→CPi raw firing rates at the time of reward (fig. S 7), decoding performance was negligible following subtraction of the mean PSTH for each reward offer (Fig. 4C,D), indicating that decoder performance in this epoch was driven by the projection neurons’ responses to rewards.

These results suggest a multiplexed temporal model, in which OFC→CPi neurons represent the belief (the full posterior probability distribution of the hidden state) during the delay period, and then shift to conveying high block evidence at the time of reward to update beliefs.

### OFC→CPi neurons impact OFC block encoding

We previously identified neural correlates of inferred states in OFC at the single neuron and population levels (*18*). We therefore hypothesized that optogenetically updating beliefs about inferred states by stimulating OFC→CPi neurons might disrupt representations of inferred states in OFC. To test this hypothesis, we recorded population dynamics in OFC while optogenetically stimulating the terminals of OFC→CPi projection neurons (Fig. 5A). We used a supervised dimensionality reduction method, demixed PCA (*33*), to identify latent factors that reflect encoding of specific task variables, including time relative to different task events, the offered reward volume, and the hidden reward state or block (Fig. 5B). We performed neurophysiological recordings for two successfully completed blocks without any perturbations, to estimate the demixed principal components in the absence of perturbations. While recording from the same neurons, we then performed optogenetic stimulation on 40% of trials while animals received water rewards. The first two optogenetic blocks were identical to the previous two control blocks, and then the alternation between high and low blocks resumed. We projected neural activity from both epochs including only trials which did not directly receive stimulation onto the components derived from the control blocks.

**Figure 5:**
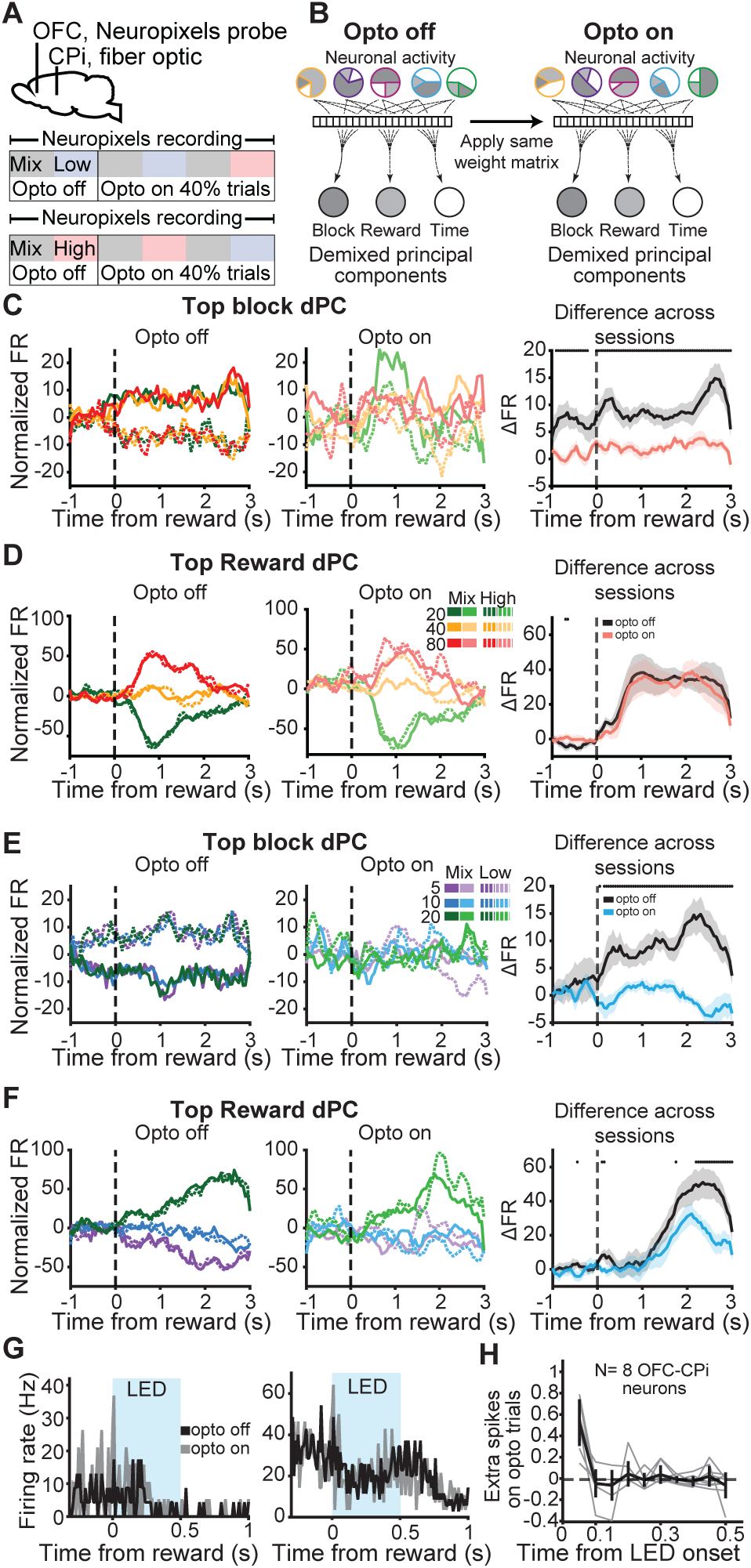
OFC→CPi perturbations impact population-level cortical representations of in-ferred states. **A.** Schematic of implantation sites (top) and task structure (bottom). To ensure we could record the activity of identical neurons with and without stimulation, LED stimulation began after two control blocks. On a given day each rat experienced a mixed block and either a high or low block with and without optogenetic stimulation. After the first four blocks, the block identity alternated normally. **B.** Schematic of demixed PCA strategy. A weight matrix was learned based on neuronal activity during trials without optogenetic stimulation, “Opto off,” to identify demixed components that explain task variables of interest: block, reward, and time. The weight matrix was applied to the same identified neurons during the optogenetic stimulation phase of the session on trials without stimulation to assess long-lasting effects on neuronal representa-tions of block and reward. **C.** PSTH of neuronal activity projected into the top block demixed principal component without (left) and with (middle) optogenetic stimulation from an example session in which the rat experienced mix and high blocks. Mean (+/- sem) change in firing rate between mix and high blocks during blocks without stimulation (black) and with stimulation (red) across all animals and sessions (N = 3 animals, n = 15 sessions) to show discriminability between mix and high block encoding. Asterisks denote a significant difference in encoding with and without stimulation (*p <* 0.05, non-parametric permutation test). **D-F.** Analyses as in C for the top reward dPC for high/mixed blocks (D), top block dPC for low/mixed blocks (E), or top re-ward dPC for low/mixed blocks (F). Actual p values for each plot are listed in Supplemental Data Tables S6-S9. **G.** PSTH of OFC→CPi neurons at reward on trials with and without LED stimu-lation. **H.** The number of extra spikes recorded from OFC→CPi somata during reward delivery on trials with LED stimulation.

Due to the requirements of implementing demixed PCA, we separately compared blocks offering large rewards in high and mixed blocks (20, 40, 80 µL), and small rewards in low and mixed blocks (5, 10, 20 µL). Projecting activity from control blocks onto the component that most strongly encoded the hidden states showed differential dynamics across blocks but not for different rewards, as expected. However, following optogenetic stimulation of OFC→CPi terminals in the striatum, representations of blocks in OFC was degraded (Fig. 5C,E). We quantified this over sessions by comparing the average difference between the projection for the different blocks (Methods). In contrast, projecting activity onto the component that most strongly encoded reward volumes showed similar encoding for both control and optogenetic blocks (Fig. 5D,F). This was especially true for large rewards, which may produce stronger inputs that make the circuit dynamics more robust to perturbations. Results were similar if we included stimulation trials in the demixed PCA analysis (fig. S 8). Therefore, stimulating OFC→CPi terminals at the time of reward disrupted population-level representations of hidden states in OFC, but largely spared encoding of rewards. We note that the reward responses identified from this analysis show that representations of value exist in OFC, consistent with prominent views of OFC function (*23*, *34*, *35*). However, those value representations do not appear to depend on the OFC→CPi neurons that update beliefs about states.

Optogenetic perturbations of axon terminals could impact OFC dynamics via back-propagating action potentials, or via long-range cortical-basal ganglia-thalamic loops. We recorded a small number (n=8) of optogenetically tagged neurons that passed collision tests in the demixed PCA experiments, which provided an opportunity to characterize the impact of perturbations that impacted behavior on the activity of the projection neurons. Surprisingly, stimulation of axon terminals produced relatively weak changes in the somatic spiking of optogenetically-tagged OFC→CPi neurons identified from collision tests, eliciting less than a single extra spike on stimulation compared to control trials (Fig. 5G,H). We speculate that these weak effects may be due to cortical inhibition that contributes to the saturating nonlinearity we previously documented at the time of reward (Fig. 3), or collisions, since this projection class was often active at the time of reward/stimulation. Therefore, OFC→CPi perturbations impact population-level cortical representations of inferred states in OFC, and based on the weak antidromic effects, we hypothesize that this modulation derives from long-range cortical-basal ganglia-thalamic loops. Future studies recording from and manipulating downstream neurons in the CPi should test this hypothesis.

## Discussion

OFC in rodents is required to infer values based on high-order associations (*13*), and to infer states with different task contingencies (*14–16*). Studies in rats and monkeys have found that OFC is important for learning when task contingencies are partially observable, potentially reflecting a key role in state inference (*14*, *17*, *36*, *37*). OFC is thought to support confidence judgments in perceptual decision-making (*38*, *39*), suggesting a role in computing or evaluating subjective belief distributions over perceptual stimuli. Our results extend these findings and show that (1) rat OFC supports belief updating (*18*), consistent with OFC playing a privileged role in learning (*27*, *40*, *41*), (2) projection-specific OFC→CPi neurons instantiate the belief updating computation, and their activation produces persistent effects that last over trials, (3) belief updating impacts OFC representations of inferred states consistent with a mechanism in which belief updates propagate through cortico-basal ganglia thalamic loops, and (4) local circuit and cellular mechanisms (potentially including inhibitory interneurons) convert graded representations of rewards into categorical representations of evidence for belief updating. The last observation suggests that OFC inhibitory interneurons may be a key locus of plasticity as animals learn to infer latent task structure from sensory features. We observed OFC dynamics reflecting value, consistent with theories that OFC represents values for neuroeconomic choices (*34*, *35*). Those dynamics were not impacted by stimulation of the OFC→CPi projection neurons, suggesting that distinct subcircuits within OFC might support these different functions.

Tasks requiring the integration of noisy sensory evidence in favor of a binary choice (*42*, *43*), or motor timing tasks (*44*), have shown that the CP integrates cortical inputs over time on single trials until a bound or threshold is reached, and a movement is initiated indicating the subject’s decision. For evidence accumulation, sensory evidence favors one of two mutually exclusive motor acts: saccade in the direction of moving dots, orient to the side with more auditory clicks. In these cases, it is conceptually straightforward that different CP neurons might support different movements, and integrate sensory inputs in favor of those movements (*45*).

While we also find that the CP is the integrator in our task, our results differ from this previous work in several key respects. First, we find that the CP integrates block evidence across trials, as opposed to within them, raising the intriguing question of what neural mechanisms can support integration on across-trial timescales. Second, evidence is integrated to compute an abstract belief about latent states, as opposed to one of two mutually exclusive motor acts. Because the reward and opt-out ports randomly change on each trial in our task, striatal neurons cannot support integration in favor of a particular left/right choice or orienting movement. Instead, they appear to integrate block evidence to compute subjective beliefs about latent states. The behavioral policy (how long to wait on each trial), is then conditioned on the reward offer, subjective beliefs about states, and the side of the reward and opt-out ports on that trial. How this is implemented at the circuit level is an important future direction.

Uncertainty in sensory and motor domains has been studied for continuous variables, and experimental evidence has supported various neural coding schemes in these domains, including probabilistic population codes (*46–49*), distributed distributional codes (*50*), and neural sampling codes (*51–54*). However, none of these traditional coding schemes are applicable to dynamic inference of categorical (as opposed to continuous) variables. While previous studies have shown that dopamine neurons can reflect uncertainty about binary categorical variables (*55*, *56*), the simple nature of this problem makes it impossible to dissociate representation of the mean from that of the full distribution. Our work is consistent with studies showing that uncertainty is represented in higher order cortical regions (*39*, *57–59*), but provides novel insight into how specific projection classes explicitly represent probability distributions over discrete, categorical variables for dynamic inference of latent states. These representations were dynamic, as OFC→CPi neurons reflected the belief distribution at trial start and as rats waited for rewards, but not at the time of reward, when they encoded high block evidence. In contrast, the CPr may not be behaviorally engaged by this task, as we did not identify a consistent behavioral effect from perturbing OFC→CPr neurons, or specific task variables encoded by that population.

At the time of reward, it is difficult to resolve the exact probabilistic variables that are being represented. The likelihood of encountering the three large rewards (20, 40, 80 µL) in a high block is equal, but the OFC→CPi neurons only encode 40 and 80 µL categorically, and have lower average firing rates for 20 µL. This could be consistent with representing the log odds of being in a high block versus not (*60*, *61*), which would yield a different response for 20 µL, but the exact mapping between the latent computation and the neural responses is difficult to resolve. Similarly, one might imagine a distributional scheme, where the positive d’ OFC→CPi neurons represent the likelihood of being in a high block and the negative OFC→CPi neurons represent the likelihood of being in a low block, such that the population represents the full likelihood distribution. However, this seems inconsistent with the results of the optogenetics experiment, which biased rats’ beliefs specifically towards high blocks. We interpret the OFC→CPi neurons as encoding evidence for high blocks, but refrain from making strong claims about whether or not they represent the likelihood *per se*.

The recent proliferation of large-scale recording methods in neuroscience has led to a paradigm shift that emphasizes the analysis of neural populations over the activity of single neurons (*62*, *63*). We highlight two points raised by our findings. First, most cognitive tasks are complex and involve multiple algorithmic computations (e.g., working memory, state inference, deliberation between alternatives). In the absence of perturbations, which computation is actually supported by the piece of tissue under study is often unclear. In our recent study, persistent inactivation of OFC with muscimol impaired rats’ ability to update their beliefs about the blocks (*18*). This is consistent with our present findings, in which transiently stimulating OFC→CPi neurons did update rats’ beliefs, but in a biased, directional way (towards high blocks), due to the tuning of the OFC→CPi projection neurons, which on average have higher firing rates for high block evidence. Perturbations were critical for generating the hypothesis that OFC→CPi neurons instantiate the belief updating computation. It is not clear that we would have inferred this function from recordings alone.

Second, while population decoding approaches implicitly assume that downstream circuits can access all neurons (or some random subset of them), it is unclear if downstream circuits uniformly sample from the population, or receive a circumscribed subset of response types. We found that OFC→CPi neurons encode specific task variables, and downstream circuits in the CPi do not uniformly sample from the entire population, consistent with past studies of OFC projec-tions to the striatum (*23*). While the overall OFC population may have exhibited heterogeneous and mixed encoding of task variables, downstream neurons in the CPi received largely “unmixed” information about block evidence at the time of reward. Our findings show that the responses of single neurons can be highly informative of their function, especially in the context of an algo-rithmic hypothesis grounded in causal perturbations.

## Acknowledgments

We thank Roozbeh Kiani and members of our labs for comments on the manuscript and helpful discussions. We thank research technicians in the Constantinople lab for animal training.

## Author Contributions

Conceptualization: MLD, CS, CMC

Methodology: MLD, DH, HG

Investigation: MLD, JS, MAW, DH, HG, SSS

Visualization: MLD, DH, HG

Funding acquisition: MLD, CS, CMC

Project administration: CS, CMC

Supervision: CS, CMC

Writing – original draft: MLD, CMC

Writing – review and editing: MLD, DH, HG, CS, CMC

## Data, code, and materials availability

The data generated in this study will be deposited in a Zenodo database upon publication. Code used to analyze all data and generate figures will be available at https://github. com/constantinoplelab/published/tree/main upon publication.

### Supplementary materials

### Materials and Methods

#### Rats

593 Long-evans rats (*Rattus norvegicus*) between the ages of 6 and 24 months were used in this study (367 males, 226 females). Rats were water restricted to motivate them to perform the task. At the end of the day, rats were given 20 minutes of access to ad libitum water. Following behavioral sessions on Friday until mid-day Sunday, rats received ad libitum water. Animal use procedures were approved by the New York University Animal Welfare Committee (UAWC #2021-1120) and carried out in accordance with National Institute of Health standards.

#### Behavior training and task logic

Rats were trained in incremental stages of task complexity in a high-throughput behavioral facility using a computerized training protocol as described previously (*8*).

LED illumination from the central nose port indicated that the animal could initiate a trial by poking its nose in that port. Upon trial initiation the center LED turned off. While in the central nose port, rats needed to maintain center fixation for a duration drawn uniformly from [0.8, 1.2] seconds. During the fixation period (“Offer cue”), a tone played from both speakers, the frequency of which indicated the volume of the offered water reward for that trial [1, 2, 4, 8, 16 kHz, indicating 5, 10, 20, 40, 80 µL rewards for male rats and 4, 8, 16, 32, 64 µL for female rats]. Female rats consumed less water due to their spaller body size and performed substantially fewer trials than the males when offered the same rewards as the males. For analysis, the reward volumes were treated as equivalent to the corresponding values for the male rats. Following the fixation period, one of the two side LEDs was illuminated, indicating that the reward might be delivered at that port (“Reward port on”); the side was randomly chosen on each trial. This event also initiated a variable and unpredictable delay period, which was randomly drawn from an exponential distribution with mean = 2.5 seconds. The reward port LED remained illuminated for the duration of the delay period, and rats were freely behaving during this period. When reward was available, the reward port LED turned off (“Reward announced”), and rats could collect the offered reward by nose poking in that port (“Reward delivered”). The rat could also choose to terminate the trial at any time by nose poking in the opposite, un-illuminated side port (“Opt Out”), after which a new trial would immediately begin. On a proportion of trials (15-25%), the delay period would only end if the rat opted out (catch trials). If rats did not opt-out within 100 seconds on catch trials, the trial would terminate.

The trials were self-paced: after receiving their reward or opting out, rats were free to initiate another trial immediately. However, if rats terminated center fixation prematurely, they were penalized with a white noise sound and a time out penalty, typically lasting 2 seconds (“Violation”). The trial following a premature fixation break offered the rats the same amount of reward in order to dis-incentivize premature terminations for small volume offers.

We introduced semi-observable, hidden-states in the task by including uncued blocks of trials with varying reward statistics (*64*): high and low blocks, which offered the highest three or lowest three rewards respectively, were interspersed with mixed blocks, which offered all volumes. Unless otherwise noted, there was a hierarchical structure to the blocks, such that high and low blocks alternated after mixed blocks (e.g., mixed-high-mixed-low, or mixed-low-mixed-high). Each session always started with a mixed block.

In a subset of sessions used to test the impact of OFC→CPi pertubation on cortical representation of inferred states we modified the hierarchical structure to the blocks, such that the first two blocks of the session were repeated twice and then began to alternate between high and low block (e.g. mixed-high-mixed-high-mixed-low). These sessions varied day-by-day so that the animal not experience one block more often than the other.

In both cases blocks transitioned after 40 successfully completed trials. Because rats prematurely broke fixation on a subset of trials, in practice, blocks varied in number of trials.

Rats were considered to have achieved task proficiency when their time to opt-out exhibited linear sensitivity to the offered reward volume and time to opt-out for the 20 µL offer in low blocks was longer than in high blocks for at least 3 consecutive days. Rats that satisfied these criteria underwent surgeries specific to experiments. After recovering from surgeries, rats were retrained until they achieved the same pre-surgery performance which typically took one session.

#### Behavioral model

We made use of the behavioral model previously established to describe rats’ behavior in the temporal wagering task (*8*, *18*). In brief, the model is adapted from Lak et al. (*38*) which described wait time (WT) in terms of the value of the environment (or opportunity cost of continuing to wait for the reward to arrive), the delay distribution, and the catch trial probability. Given an exponential delay distribution we predict the wait time as:

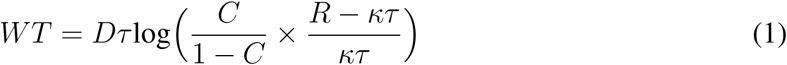

where *τ* is the time constant of the exponential delay distribution, *C* is the the probability of reward, *R* is the reward volume on the current trial, *κ* is the opportunity cost, and *D* is a scaling parameter. The decision to opt out depends on the opportunity cost, *κ*, which is stable within trials but can vary across trials and hidden reward states, i.e. blocks. The inferential model has three discrete opportunity cost parameters associated with each block (*κ_low_*, *κ_mix_*, *κ_high_*). On each trial, the model chose the *κ* associated with the most probable block given the recent reward history according to Baye’s theorem, described by:

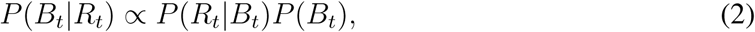

where *B_t_*is the block and *R_t_*is the reward on trial *t*. The likelihood, *P* (*R_t_*|*B_t_*) is the probability of the reward for each block. For example:

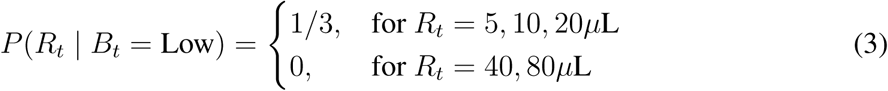

To calculate the prior, *P* (*B_t_*), we marginalized over the previous block and used the previous estimate of the posterior:

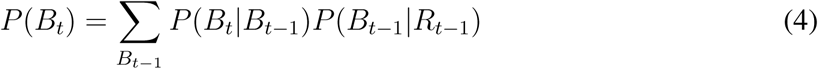

The “hazard rate,” *P* (*B_t_*|*B_t_*_−1_) incorporates knowledge of the task structure, including the block length and block transition probabilities. For example:

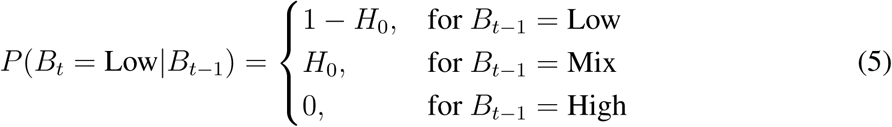

where *H*_0_ = 1/40 to reflect the block length. Including *H*_0_ as an additional free parameter has been shown previously to not improve the performance of the inferential model on held-out test data (*8*), so *H*_0_ was treated as a constant term. See (*8*) for additional discussion of the hazard rate.

#### Biased behavioral model

For the model with a biased prior, we included a term that was added to the probability of being in a high block, and subtracted from the low and mixed block probabilities as in:

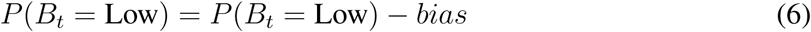

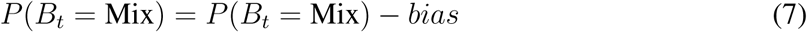

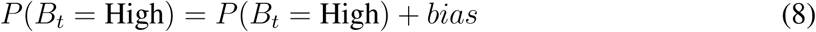

After the additive bias was introduced, negative beliefs were set to zero and the block probabilities were normalized to values between zero and one. The bias used for the model simulations was set to 0.04.

In separate models we modulated the opportunity cost of all blocks by increasing the *κ* for each block by 0.15, or modulated the time constant of the time-varying value function, *τ*, by reducing its value from 2.5 to 2.0. In a third model we replaced the offered reward, *R_t_*, with 80 *µ*L on 40% of trials, to simulate a perturbation that altered the perception of reward volume.

### Surgery

Rats were anaesthetized (isofluorane at 3% for induction and 1.5-2.5% for maintenance) and placed in a stereotaxic frame. In experiments in which hardware was attached to the skull, skull screws were inserted to secure implants before proceeding with surgeries specific to experiments. Target coordinates relative to the skull surface at bregma for the OFC were: AP: 3.7 mm, ML: ± 2.9mm, DV: 5 mm, the CPi: AP: -0.2 mm, ML: ± 3.5 mm, DV: 4.5 mm, and CPr: AP: 1.2 mm, ML: ± 2.5 mm, DV: 6.9 mm, SNr: AP: -5.0, -5.3, -5.7 mm, ML: ± 2.3 mm, DV: 8.4, 8.3, 8.2 mm, and GPe: AP: -1.5 mm, ML: ± 3.3 mm, DV: 6.8 mm.

#### Anatomical experiments

For anterograde tracing of OFC projection neurons, rats were injected intracranially with 100 nL of pENN.AAV.CB7.CI.mCherry.WPRE.RBG (AAV9) (AddGene, #105544-AAV9) in the OFC. Viral expression was given 2 weeks to mature before animals were sacrificed.

For retrograde tracing of OFC projection neurons, rats were injected with 120 nL of Alexa fluorophore 657 conjugated cholera toxin subunit B (ThermoFisher Scientific, C34778) in either the CPi or CPr. CTB was diluted at 0.5% (wt/vol) in PBS (*65*). Following the injection, the burr hole with filled with Kwik-Sil (WPI), and the skin was sutured. The CTB was given 10 days to spread retrogradely to somata.

#### Optogenetic stimulation experiments

For optogenetic manipulations and optotagging protocols, rats were injected with 500 nL of retrograde-serotype Cre virus (“Cre_rg_”, AAVrg-pgk-Cre, AddGene #24593) in the CPi or CPr, and Cre-dependent channelrhodopsin (“ChR2”, AAV9-EF1a-double floxed-hChR2, AddGene #20298) in the OFC. 100 nL of virus was injected at a time at a rate of 10 nL sec^−1^.

Rats used for optogenetic manipulation, but not electrophysiology, were then immediately bilaterally implanted with a tapered fiber optic (0.66 NA, 1 mm active length, 200 µM core, Optogenix) in the CPi or CPr such that the emitting tip of the fiber was within the target region. The fiber was secured to the skull surface using dental cement and Dentin and then the headcap was sealed with dental acrylic. Rats were allowed to recover for a minimum of 5 days. Rats performed the task for at least 2 weeks before any LED stimulation was delivered during the task to allow for cells to fully express the virus.

#### Electrophysiological experiments

Rats used for optotagged electrophysiological experiments underwent two distinct surgeries; one to inject viruses and a second to implant physiological hardware after the viruses were fully expressed. We found that chronically implanted Neuropixels probes could be damaged or signals could degrade sufficiently before the ChR2 virus was fully expressed. Following viral injections as described for optogenetic manipulation, all injection site burr holes were sealed with Kwik-sil (WPI) and the skin sutured. Viruses were given 2-6 weeks to express. During this period rats were first given 5 days to recover, and then were returned to performing the task daily to ensure their behavior remained stable.

The day before the implantation surgery, a Neuropixels probe assembly was prepared by mounting the probe (Neuropixels probe 1.0 or 2.0) onto a custom-designed 3D printed piece based on other chronically implanted headcaps (*66*) and staining with DiI. Open Ephys built-in self tests were run on the probe assembly prior to surgery to ensure all probe shanks were functional prior to implantation.

During the second surgery, fiber optics were implanted bilaterally in the CPi or CPr, and the Neuropixels probe was implant using the probe assembly in the OFC. A silver ground wire that shorted the ground and reference connection on the probe was inserted in the cerebellum and fixed with Dentin (Parkell). Craniotomies were covered with Kwik-Cast (WPI) and gaps between the probe assembly and the skull were sealed with sterile petroleum jelly. The probe and fibers were then fixed to the skull with Metabond (Parkell) and Absolute Dentin (Parkell), and sealed with Ortho-jet dental acrylic (Patterson Dental Comparny). Bubble wrap was placed around the implant to protect it from damage. Rats were allowed to recover for a minimum of 5 days before returning to performing the task and electrophysiological recording.

### Histology

Following completion of each experiment, rats were deeply anesthetized and transcardially perfused with phosphate buffer saline (PBS) and formalin. Brains were post-fixed with formalin for 3 days and sectioned at 50 *µ*m slices on a Leica VT1000 vibratome.

#### Immunohistochemistry

To confirm ChR2 expression, ChR2-conjugated EYFP signal was amplified using antibody labeling. Sections were treated with 1% hydrogen peroxide in 0.01 M PBS for 30 min, washed with 0.01 M PBS, and blocked in 0.01 M PBS containing 0.05% sodium azide and 1% bovine serum albumin (PBS-BSA-Azide) for 30 minutes prior to treating with primary antibodies. EFPF was amplified with Rabbit anti-GFP (Thermo Fisher Scientific A11122, 1:2,000, lot #2083201) as primary antibody and Goat anti-Rabbit AlexaFluor 488 IgG (Thermo Fisher Scientific A11008, 1:200, lot #2147635) as secondary antibody. Primary and secondary antibodies were made in PBS-BSA-Azide and primary antibodies additionally contained 0.2% Triton-x. Sections were incubated in primary antibody overnight at room temperature, washed with PBS for 15 min, and treated with secondary antibody for 1 hour in the dark. After washing with PBS for 15 min, sections were mounted on glass slides using Prolong Gold (Vector Laboratories).

#### Imaging

For electrophysiology and optogenetic experiments, sections were imaged with an Olympus VS120 Virtual Slide Microscope to verify probe and fiber optic locations, and ChR2 expression. Neuropixels probe tracks were reconstructed from post-mortem histology, and the location of individual recording channels relative to areal boundaries was estimated. Channels that were estimated to be outside of OFC or agranular insula (AI) were excluded from further analysis.

For anatomical experiments, injection sites were confirmed using 10x magnification on an Olympus VS120 Virtual Slide Microscope. Light intensity was controlled across all sections. Cell bodies were imaged at 10x and 20x magnificantion using a Leica DM6 CS Confocal with standardized parameters developed to maximize signal-to-noise ratios. For each animal 3 sections near the target location of AP: Bregma +3.7 mm were selected by anatomical markers and imaged. Brain sections were then manually aligned to standardized brain sections described in Paxinos et. al. (*67*) to reduce variability caused by brain size, or stretching/shrinking during processing or mounting, and enable averaging across animals. OFC projection neurons somata were manually labeled and counted using the ImageJ cell counting pluggin which tracks number and location. Cell number was averaged within 250 *µ*m^2^ bins tiling the brain section and then further averaged across the 3 sections imaged for each animal. We then calculated the mean cell number in each tile across animals.

### Electrophysiological recordings

#### Neuropixels data acquisition

Spike data was acquired from 384 channels at 30 kHz using Neuropix-PXI hardware and OpenEphys (https://github.com/open-ephys). To synchronize physiological recordings with behavior and optogenetic LED pulses, TTL pulses were recorded simultaneously using NI-DAQmx. NI-DAQmx and Neuropix-PXI recordings were aligned using a six digit temporal barcode generated by an Arduino Zero and simultaneously sent to both recordings.

#### Spike sorting

Spike data were preprocessed using Kilosort 2.5. After preprocessing, clusters that were identified from Kilosort as single-units were manually inspected using the Phy2 Python pack-age (https://github.com/cortex-lab/phy, no longer maintained). Clusters identified by Kilosort as “good” units with unreliable waveform shape, autocorrelogram, or spike amplitude time course were manually classified as multi-unit or noise and removed from further analysis. Units were further curated using a custom MATLAB script. Units with greater than 1% inter-spike intervals less than 1 ms, firing rates less than 1 Hz, or completely silent for more than 5% of the total recording were excluded. To convert spikes to firing rates, spike counts were binned in 50 or 200 ms bins where described in the text.

#### Identification of fast-spiking interneurons

After manual curation, we used the template waveforms generated from Kilosort 2.5 to measure the half-width spike duration. Specifically, we used the time from the minimum waveform trough to the following largest peak. We found that neuron in our OFC recording fell into a bi-modal distribution, and we used the time point of the minumum between the two peaks (0.4 ms) as a cut off between regular-spiking neurons and fast-spiking interneurons.

#### Optogenetic identification of projection specific neurons

Optogenetic tagging of projection specific neurons was performed after the rat had completed its 90 minute behavioral session to prevent optogenetic stimulation influencing the rat’s behavior. Blue light (465-nm PlexBright LEDs, Plexon) was delivered in 20 trains of 10 pulses at 10 Hz, 3 ms. During the optotagging protocol rats were left in the behavioural chamber without a task to perform so that stimulation was unlikely to affect task performance on subsequent days. Optical power emited from the patch cable was measured daily prior to the animal performing the task and the current applied to the LED was modulated to maintain an average of 10 mW power during a 1 s continuous pulse using a Thorlabs optical Power and energy meter (PM100D2).

Neurons which reliably generated at least one action potential (i.e. with response to a greater number of LED pulses than two standard deviations from the mean of all OFC cells) within the same 2 ms window following LED onset and with a latency less than 20 ms were further assessed for action potential collisions. These criteria reduced the number of cells assessed for collisions from 7697 to 251 cells. We manually assessed LED pulse ‘trials’ in which the neuron elicited an action potential before the median latency of the first action potential after LED onset. If a spike occurred before the median latency, as well as at the expected delay following LED onset, the neuron failed the collision test, and was not included as a direct projection neuron.

### Optogenetics experiment

After recovering from surgery, rats were habituated to being tethered until they achieved pre-surgery performance which typically took 1-2 sessions. Following habituation, control sessions (no LED) and stimulation sessions (LED on on 40% of trials) were alternated every 5 days for a minimum of 12 sessions total per rat. A subset (3) animals received electrophysiological implants to determine the impact of OFC→CPi perturbations on OFC activity directly. These animals underwent two surgeries as described for electrophysiological implants, so were fully expressing ChR2 at the time of implant and underwent a modified task as described for demixed PCA. As the neuropixels probes failed, animals were transitioned back to whole sessions of either control (no LED) or stimulation (LED on on 40% of trials). Blue LED modules (465 nm, PlexBright LEDs, Plexon) were connected to a commutator and 200 µM core patch cords that were tethered to the fiber implants of rats. The connections between the LEDs, patch cords, and fiber implants were sealed with black electric tape or black shrink tubes to block light leak. Continuous blue light (465-nm PlexBright LEDs, Plexon) was delivered randomly on 40% of trials for 500 ms during reward delivery. Optical power was measured daily prior to the animal performing the task and the current applied to the LED was modulated to maintain an average of 10 mW power during a 1 s continuous pulse using a Thorlabs optical Power and energy meter (PM100D2). We found that the same LED current settings during a 500ms pulse resulted in an measured average power output of 1.2-2.3 mW, although this likely reflects the capability of the light meter rather than the LED itself.

### Quantification and statisical analyses

Exact p-values were reported if greater than 10^−20^. For p-values smaller than 10^−20^, we reported *p <<* 0.001

#### Detrending wait times

To account for modest, gradual increases in wait times over the course of the behavioral session, and remove potential effects of motivation and satiety from the wait times, we performed the following detrending procedure for all wait time analyses. First, we removed outlier wait times that were one standard deviation above the pooled-session mean. We fit a regression line to the mean wait time as a function of trial number, averaging over all sessions, using MATLAB’s regress.m function. We then subtracted the regression line from the wait times on each session, adding back the offset parameter so that wait time units were in seconds and not zero-meaned.

#### Wait time sensitivity to reward blocks

For all analyses, we removed wait times that were one standard deviation above the pooled-session mean. Without thresholding, the contextual effects are qualitatively similar, but the wait time curves are shifted upwards because of outliers that likely reflect inattention or task disengagement. When assessing whether a rat’s wait time differed by blocks, we compared each rat’s wait time on catch trials offering 20 *µ*L in high and low blocks using a non-parametric Wilcoxon rank-sum test, given that the wait times are roughly lognormally distributed.

To determine the effect of optogenetic stimulation on wait times, we normalized the rat’s wait time to their mean wait time for 20 *µ*L trials in control session mixed blocks. Because each animal underwent both control and optogenetic sessions, we used Wilcoxon signed-rank test to quantify change in wait time with optogenetic manipulations. To measure difference across the entire block caused by optogenetic stimulation, we averaged the normalized wait times for all volumes in each block, and subtracted the mean wait time for each block in control sessions from the mean wait time for each block in optogenetic stimulation sessions. We used a Friedman test to assess for difference across all blocks, and when appropriate used Wilcoxon signed-rank tests to test individual comparisons.

#### Block transition dynamics

To assess behavioral dynamics around block transitions, for each rat, we detrended wait times over the course of the session, and then z-scored wait-times for all catch trials. We then z-scored wait times for each volume in each time bin (trial relative to block transition), before averaging over all volumes. We smoothed the average curve for each rat using a 5-point causal filter, before averaging over rats.

Because rats receiving optogenetic stimulation had a limited number of trials, we did not always have wait times for each volume at each time bin relative to block transition, so we combined trials into 5 or 10 trial windows around block transitions. We then compared wait times within these windows in control and optogenetic sessions using a Friedman test, and Wilcoxon signed-rank test for individual comparisons when appropriate.

#### Discriminability index

To measure encoding discriminability used a d’ metric which used the following equation:

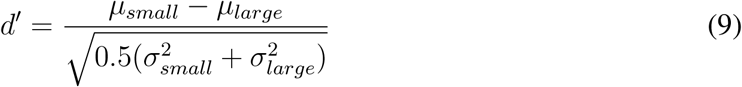

where *µ_small_* represented the mean firing rate on smaller reward volume mixed block trials and *µ_large_* represented the mean firing rate on larger reward volume mixed block trials, and 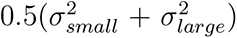 is the mean variance. For unsigned d’ values we used the absolute value of the mean difference instead. We compared individual neurons d’ values at each time point to a label-shuffled control. Significant d’ during reward delivery for individual neurons was defined as values which exceeded (positively or negative) the 95th percentile of the label-shuffled control for the entire [0 1s]. To compare d’ across different projection-specific populations, we subtracted the mean d’ of the label-shuffled control. Shuffles were computed 100 times and maintained a balance of low and high reward trial, and used the 200 ms time bins to generate the shuffle, as this was the bin-width used for computing the reward-aligned firing rates. We indicate d’ values greater than the 95th percentile of the label-shuffled d’ for that population with a larger line width in each figure to indicate that the mean d’ was signficantly different from the label-shuffled control. We then only compared d’ values between different projection classes at time points where the projection specific neurons’ d’ was greater than the 95th percentile of the label-shuffled d’ for that population. We denote time points at which the d’ of OFC→CPi or OFC→CPr differed from the untagged OFC population using a nonparametric permutation test with lines above the data.

#### Decoding reward volume

We used firing rates from different projection classes to decode reward volume on each trial. We decoded low volumes (5/10 *µ*L) or high volumes (40/80 *µ*L) on mixed block trials using neural data in 6 ms time bins. We trained logistic regression decoders (SciKit-Learn) using a 2-fold cross validation where within each fold a decoder was fit to a training set using a no penalty, saga solver. We report the average performance for decoding held-out test data for each fold.

#### Decoding most likely and second most likely block

We used neural responses from optotagged OFC→CPi neurons to separately classify each trial’s most likely block and second most likely block on each trial. These class labels were generated from a Bayes optimal model’s estimation of block probabilities (*8*). We only decoded from rewarded trials, and used neural data averaged over 200 ms time bins as the regressors. We took a pseudo-population approach: data across multiple sessions from non-simultaneously recorded neurons was used, and for each “pseudo trial” that was decoded, we chose a trial each from neuron’s recordings that shared the same class label.

The data sampling procedure was as follows: we performed 20-fold cross validation, where for each fold we randomly sampled neurons (with replacement) with correct class labels to obtain pseudo populations for 100 pseudo trials as a test set for each condition (i.e., 100 low blocks, 100 mixed blocks, 100 high blocks). For the test set, we generated those 100 pseudo trials from 20 distinct trials for each condition (i.e, 20 distinct low block trials, 20 distinct mixed block trials, 20 distinct high block trials). The same procedure was performed for a training set of 500 pseudo trials for each class label, but the neural recordings from these 60 withheld trials for testing data were not used to ensure that they were never seen by the decoder. Decoding was trained with Matlab’s ‘fitmnr‘ function for every 200 ms time bins from [-1,3]s around reward delivery, and testing accuracy was averaged over the withheld test data set. This process was performed 20 times, and reported accuracies are averaged over all 20 folds. One recording session did not contain enough trials of each type to adequately build the testing data set, which forced us to omit 3 neurons from the decoding. Thus, 57 neurons were used in the decoder.

In order to build a null distribution, we shuffled the trial labels of the optotagged recordings, and performed the same analysis using 20-fold cross validation. We performed the shuffle 100 times, and reported the results as the distribution over fold-averaged test accuracies. Finally, we also performed the analysis using data from all OFC units. In order to balance the statistics of neurons used across OFC→CPi projectors and all OFC, for each of the 20-fold regressions we randomly sampled 57 neurons, and then performed the regression as above. We performed this sampling and fitting procedure 100 times. To assess if OFC→CPi significantly decoded most likely block and second most likely block more than the general OFC population, we performed rank sum tests in each 200 ms time bin, and assessed significance at *p <* 0.05. We performed two versions of this decoding analysis. The first form used the raw firing rates from each trial epoch as regressors into the decoding. The second form removed the contribution of the reward/offer signal by subtracting off the psth of each offer size, but marginalized over blocks.

Specifically, we calculated the average psth response for each volume size, but averaged over the their contributions across all blocks. We then subtracted this marginal PSTH signal from trials with that offer/reward size, and used this in the decoding analysis. When visualizing the decoding results, we applied a causal filter over 3 data points (600ms).

#### Demixed principal component analysis

To assess the impact of projection specific manipulations on population-level cortical representations of inferred states we used demixed principal component analysis (*33*) (dPCA, software available at: http://github.com/machenslab/dPCA). This aims to decompose the data into interpretable latent components that explain variance in the data. We used reward volume, block identity, and time (task event aligned temporal dynamics), as categorical predictors of neuronal activity. Because dPCA requires that each variable is present in each category we were unable to compare high and low blocks directly as high and low blocks include non-overlapping reward volumes. Therefore, we compared mixed to low blocks and mixed to high blocks for each session. To ensure the rat experienced a high/low block in both the control and stimulation phases of a single session, we modified the task design such that on alternating days the animal experienced a mixed and high or low block with the LED turned off (“Opto off”) and those two blocks repeated once the LED was turned on for 40% of trials (“Opto on”). After the first two blocks of the Opto on phase, the blocks resumed alternation between high and low blocks interspersed with mixed blocks as in the standard task. For analysis, we required that all sessions had at least 2 trials of each reward volume in each block and binned spikes into 50 ms time bins to generate firing rates.

For each session, we first generated a weight matrix and demixed principal components (dPCs) for each session using the activity during the first two blocks (mixed and low, or mixed and high) in the session during the Opto off phase. We projected neuronal activity onto the top block or reward dPCs (i.e., the block and reward PCs that explained the most variance) to generate PSTHs of activity for each category. To generate PSTHs of the neuronal activity during Opto on, we projected neuronal activity recorded during the Opto on phase using the weight matrix and principal components estimated from control trials. To assess the long-lasting influence on OFC neuronal encoding we selected specifically trials without optogenetic stimulation (Fig. 5c-f). However, we also found similar results if we additionally included trials with optogenetic stimulation (fig. S 8)

To compare across sessions, we calculated the mean difference in block or reward encoding during the Opto off and Opto on phases across all sessions. Specifically, we computed the mean difference between the preferred block (which ever had the higher firing rate) and nonpreferred block for each reward volume (20, 40, 80 *µ*L for high vs. mixed block comparisons, and 5, 10, 20 *µ*L for low vs. mixed block comparisons) in the Opto off and Opto on phase for each session and then averaged these values across all sessions. To determine changes in reward encoding, we found the mean difference between the highest and lowest volumes (20 and 80 *µ*L for mixed vs. high and 5 and 20 *µ*L for mixed vs. low) in the Opto off and Opto on phase for each session and then averaged these values across all sessions.

---

**fig. S 1:**
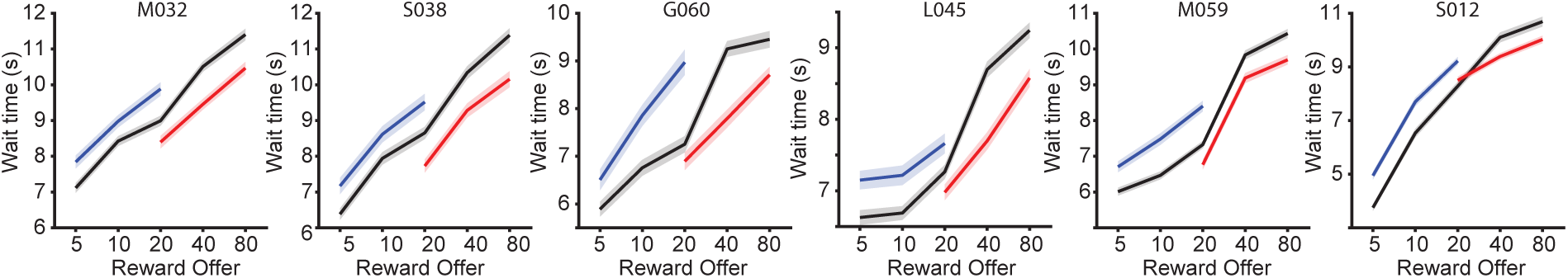
Wait time curves for individual example rats.

**fig. S 2:**
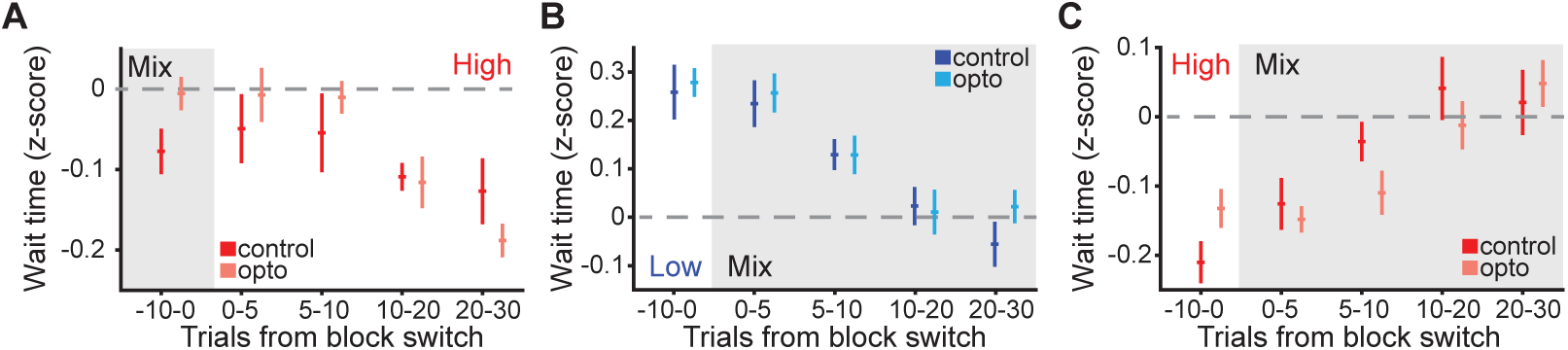
Rat wait times at other transitions are unaffected by stimulation. **A.** Mean wait time across all animals in control and opto sessions around mixed to high block transitions grouped into 5 trial bins. Friedman test, *p* = 0.5867). **B.** Mean wait time across all animals in control and opto sessions around low to mix block transitions grouped into 5 trial bins. Friedman test, *p* = 0.7353). **C.** Mean wait time across all animals in control and stimulation sessions around high to mix block transitions grouped into 5 or 10 trial bins. Friedman test, *p* = 0.7803).

**fig. S 3:**
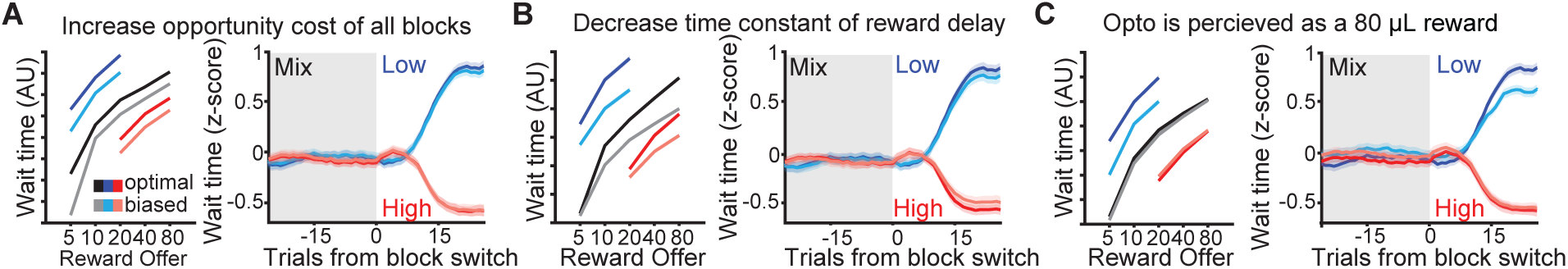
Alternative model parameters do not reproduce animal behavior. **A.** Model in which the opportunity cost of all blocks was increased decreases wait times, but does not modulate transitions. **B.** Model in which the time constant of the reward delay decreases. **C.** Model in which the reward offer is instantiated as 80 µL on stimulation trials regardless of the block or actual offered reward.

**fig. S 4:**
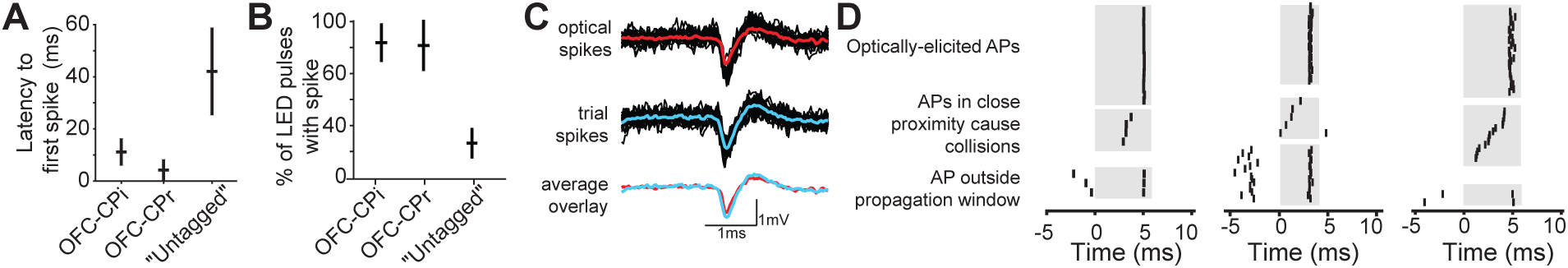
Characteristics of optically tagged projection neurons. **A.** Mean (+/- sem) latency of the first spike following LED stimulation for each OFC population class. **B.** Percentage of LED pulses with spike responses for each OFC population class. **C.** Example traces (black) and mean waveform (color) for the same neuron in response to LED stimulation (top), or elicited during behavioral trials (middle). The mean waveform of spikes both periods overlayed (bottom). **D.** Raster plots of spikes from three additional example OFC→CPi neurons which pass collision tests.

**fig. S 5:**
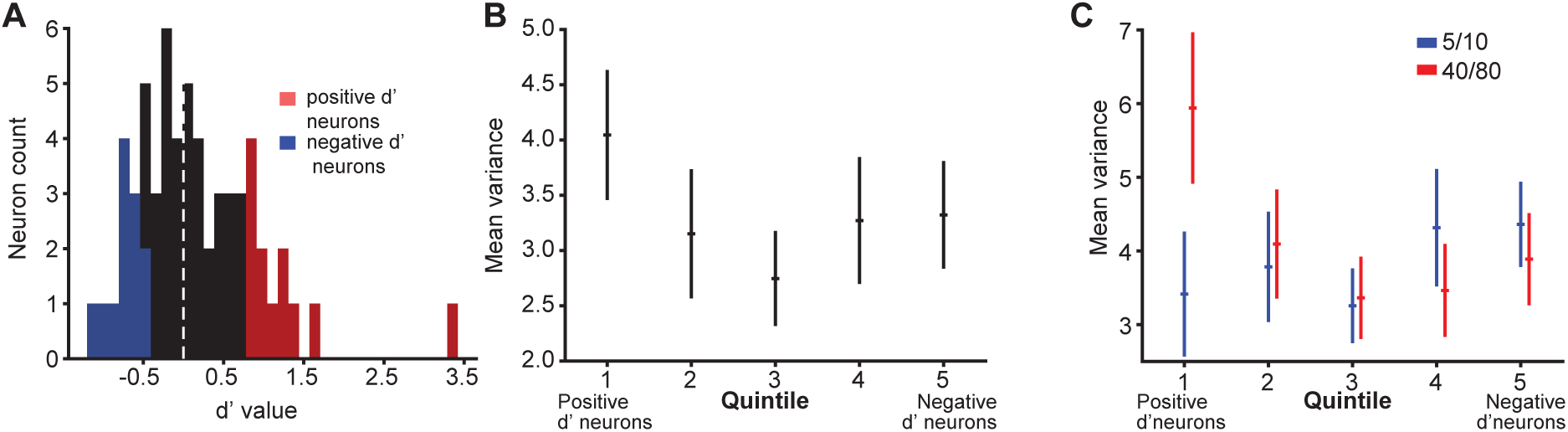
Neurons with highly positive or negative d’ do not arise from reduced within-group variance. **A.** Histogram of OFC→CPi neurons’ discriminability between 5/10 versus 40/80 µL rewards. The subset of positive and negative d’ neurons used in Fig. 3 are highlighted in red and blue respectively. **B.** Mean variance in firing rates for 5/10 and 40/80 µL rewards did not significantly differ across the population of neurons. 1-way ANOVA, *p* = 0.45097. **C.** Mean variance in firing rates between encoding of 5 and 10 µL rewards (blue) or 40 and 80 µL rewards (red) did not significantly differ across the population, within groups, or within the interaction between quintile and reward pairing. 3-way ANOVA, *p* = 0.16498.

**fig. S 6:**
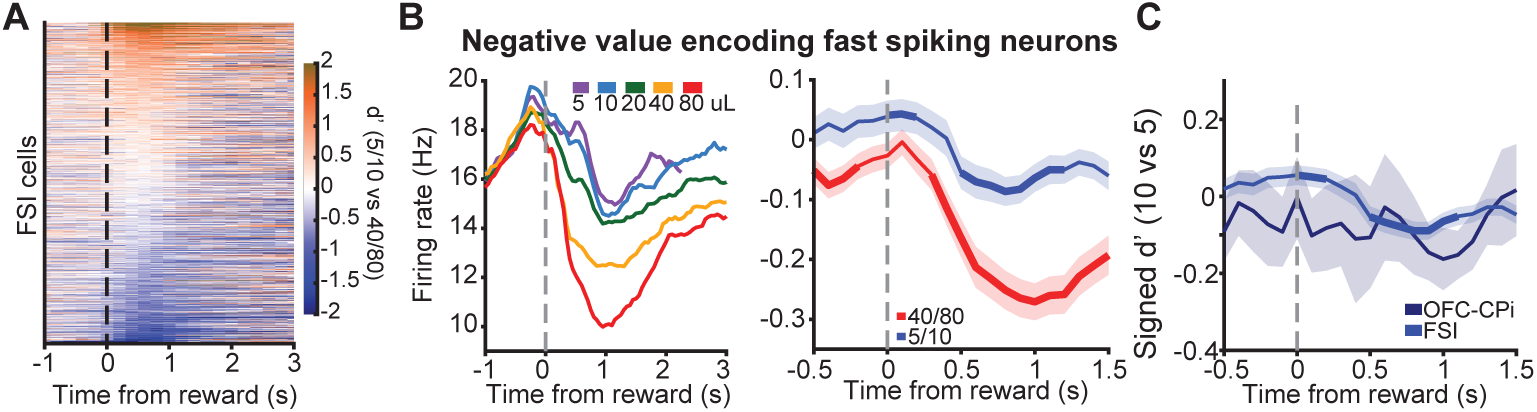
Putative fast-spiking interneurons (FSIs) show heterogenous encoding. **A.** Heatmap of signed discriminability index (d’) for trials offering 40/80 versus 5/10 µL rewards in mixed blocks for each FSI aligned to reward delivery. **B.** Negative d’ FSIs show weak discriminability between preferred rewards. Left: Mean PSTH of negative d’ FSIs for different rewards, aligned to reward delivery. Middle: Discriminability of preferred (5 versus 10 µL) and non-preferred (40 versus 80 µL) rewards. **C.** Overlay of d’ for preferred rewards for negative d’ FSIs (light blue) and negative d’ OFC→CPi neurons (dark blue).

**fig. S 7:**
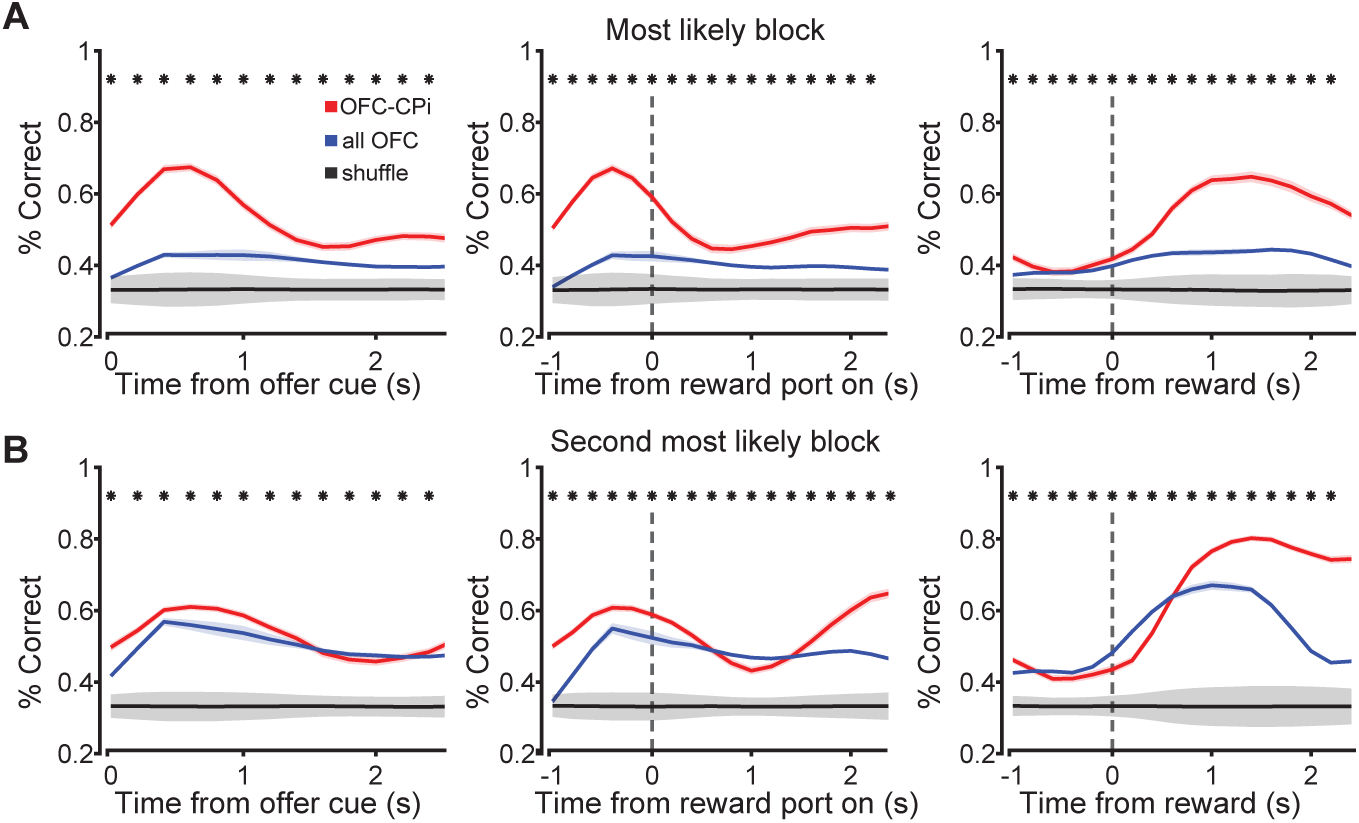
**A.** Decoding accuracy of OFC→CPi neurons to classify the most likely block, using raw firing rates (i.e., without mean-subtracting the PSTH for each reward). Colored lines represent different populations (red = OFC→CPi, blue = all OFC, black = shuffle distribution built by shuffling trials labels of OFC→CPi data. Error bars for red and blue curves are sem, and error bars for the shuffle distribution are 1.96 standard deviations calculated over 100 shuffles. Significant differences in decoding accuracy between OFC→CPi neurons and a sample-number matched untagged OFC population via Wilcoxon signed-rank test (*p <* 0.05) is denoted above each timepoint (actual p values listed in Supplemental Data Tables S3 and S4).

**fig. S 8:**
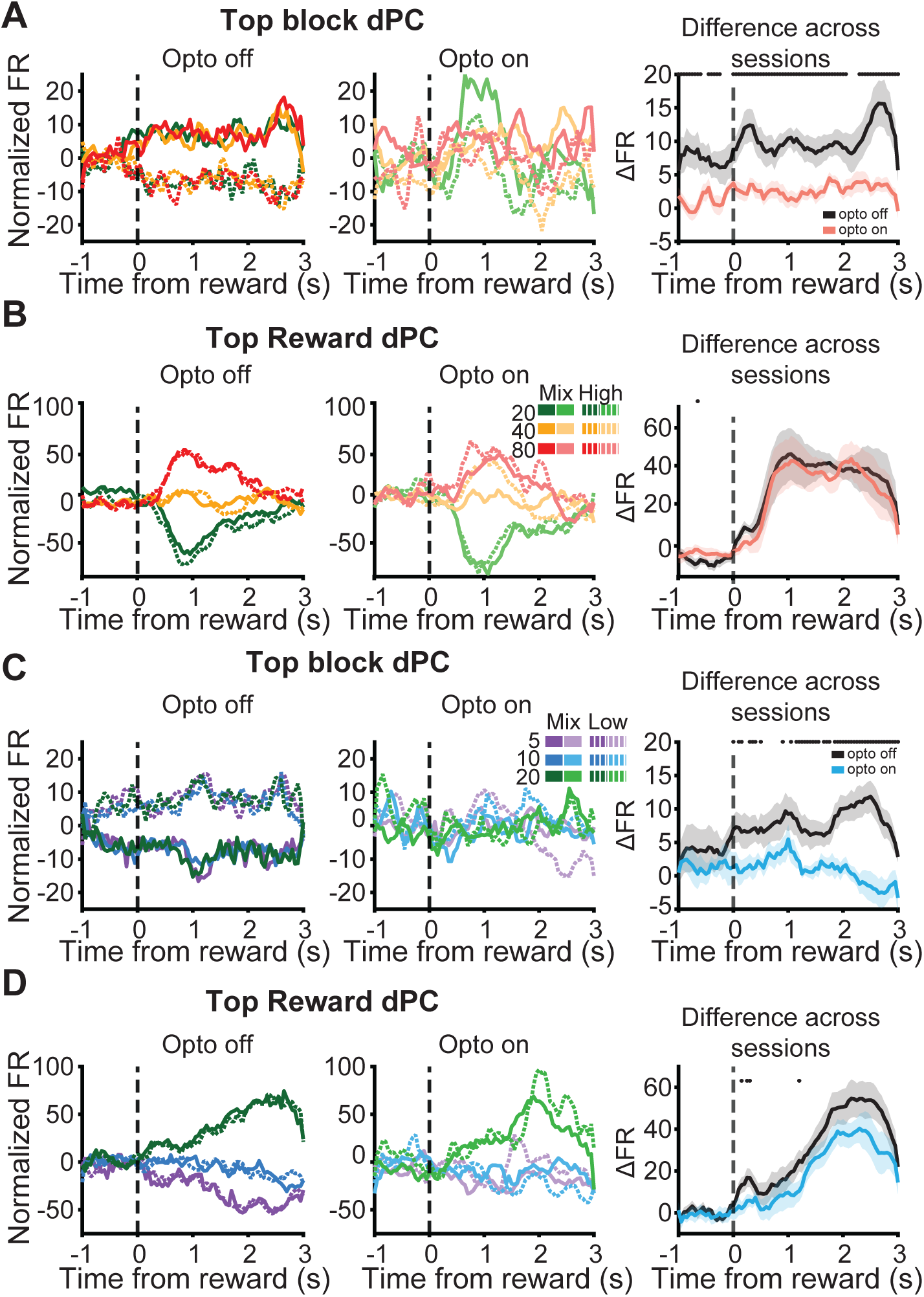
Demixed Principal Component analysis using all rewarded trials shows that opto-genetic stimulation disrupts block encoding but not reward encoding. **A.**PSTH of neuronal activity projected into the top block demixed principal component without (left) and with (middle) optogenetic stimulation from an example session in which the rat experienced mix and high blocks. Right: mean (+/- sem) change in firing rate between mix and high blocks during blocks without stimulation (black) and with stimulation (red) across all animals and sessions (N = 3 animals, n = 15 sessions) to show discriminability between mix and high block encoding. Asterisks denote a significant difference in encoding with and without stimulation (*p <* 0.05, non-parametric permutation test). **B-D.** Analyses as in **A** for the top reward dPC for high/mixed blocks (B), top block dPC for low/mixed blocks (C), or top reward dPC for low/mixed blocks (D). Actual p values for each plot are listed in Supplemental Data Tables S10-S13.

## Supplemental Data

**Supplemental Data Table S1.** The number of optically-tagged cells per rat for each recording session.

**Supplemental Data Table S2.** Non-parametric permutation test p values for each time point comparing the discriminability of large and small reward volumes between untagged OFC population and OFC→CPi neurons and OFC→CPr neurons.

**Supplemental Data Table S3.** Wilcoxon signed-rank test p values for each time point comparing decoding accuracy of the most likely block from OFC→CPi neurons to the OFC population.

**Supplemental Data Table S4.** Wilcoxon signed-rank test p values for each time point comparing decoding accuracy of the second most likely block from OFC→CPi neurons to the OFC population.

**Supplemental Data Table S5.** Non-parametric permutation test p values for each time point comparing encoding of the second most likely block on 20 µL trials when the probability of the trial being a mix block is high.

**Supplemental Data Table S6.** Non-parametric permutation test p values at each time point to compare encoding of high and mix blocks during the control and optogenetic stimulation epochs of the session, specifically using trials optogenetic stimulation was not delivered.

**Supplemental Data Table S7.** Non-parametric permutation test p values for each time point to compare reward encoding between 20 and 80 µLduring the control and optogenetic stimulation epochs of the session, specifically using trials optogenetic stimulation was not delivered.

**Supplemental Data Table S8.** Non-parametric permutation test p values for each time point comparing encoding of low and mix blocks during the control and optogenetic stimulation epochs of the session, specifically using trials optogenetic stimulation was not delivered.

**Supplemental Data Table S9.** Non-parametric permutation test p values for each time point comparing reward encoding between 5 and 20 µL during the control and optogenetic stimulation epochs of the session, specifically using trials optogenetic stimulation was not delivered.

**Supplemental Data Table S10.** Non-parametric permutation test p values for each time point encoding of high and mix blocks during the control and optogenetic stimulation epochs of the session, using all trials in epoch which includes trials in which optogenetic stimulation was delivered.

**Supplemental Data Table S11.** Non-parametric permutation test p values for each time point to compare reward encoding between 20 and 80 µL during the control and optogenetic stimulation epochs of the session, using all trials in epoch which includes trials in which optogenetic stimulation was delivered.

**Supplemental Data Table S12.** Non-parametric permutation test p values for each time point comparing encoding of low and mix blocks during the control and optogenetic stimulation epochs of the session, using all trials in epoch which includes trials in which optogenetic stimulation was delivered.

**Supplemental Data Table S13.** Non-parametric permutation test p values for each time point comparing reward encoding between 5 and 20 µL during the control and optogenetic stimulation epochs of the session, using all trials in epoch which includes trials in which optogenetic stimu-lation was delivered.

---

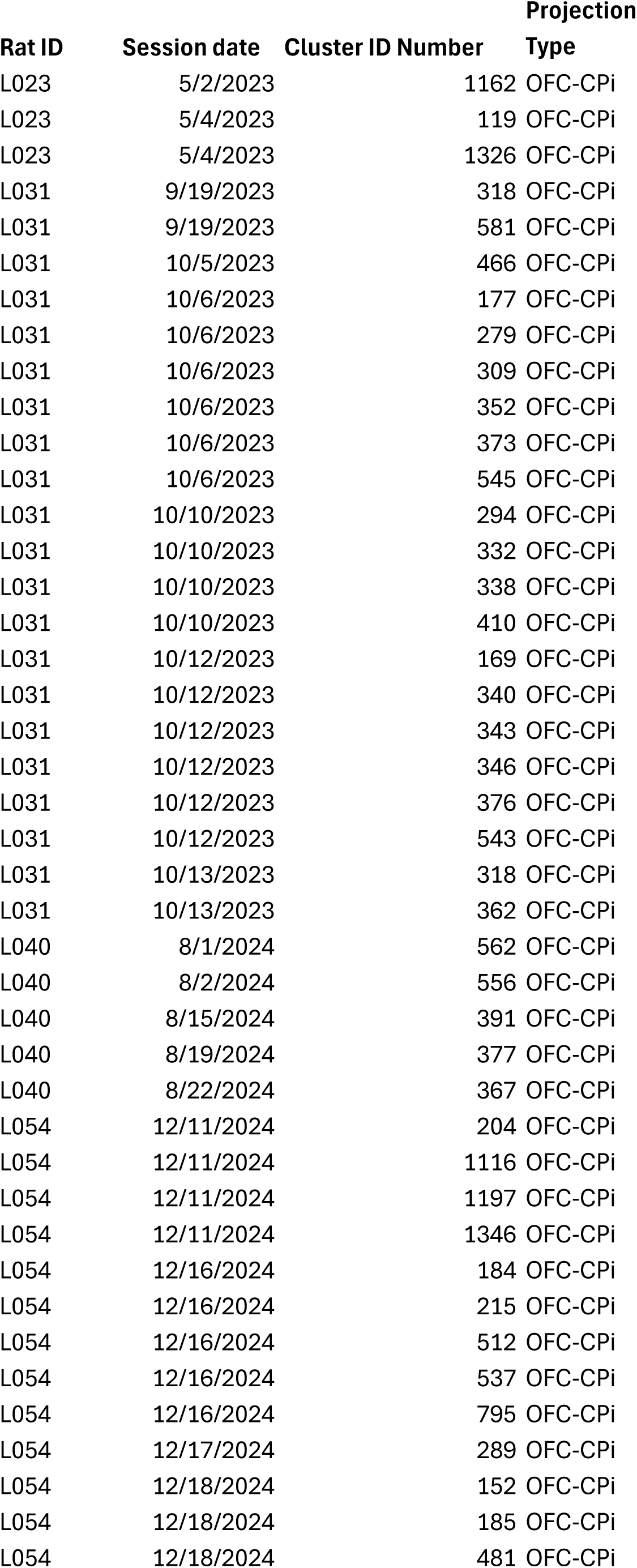
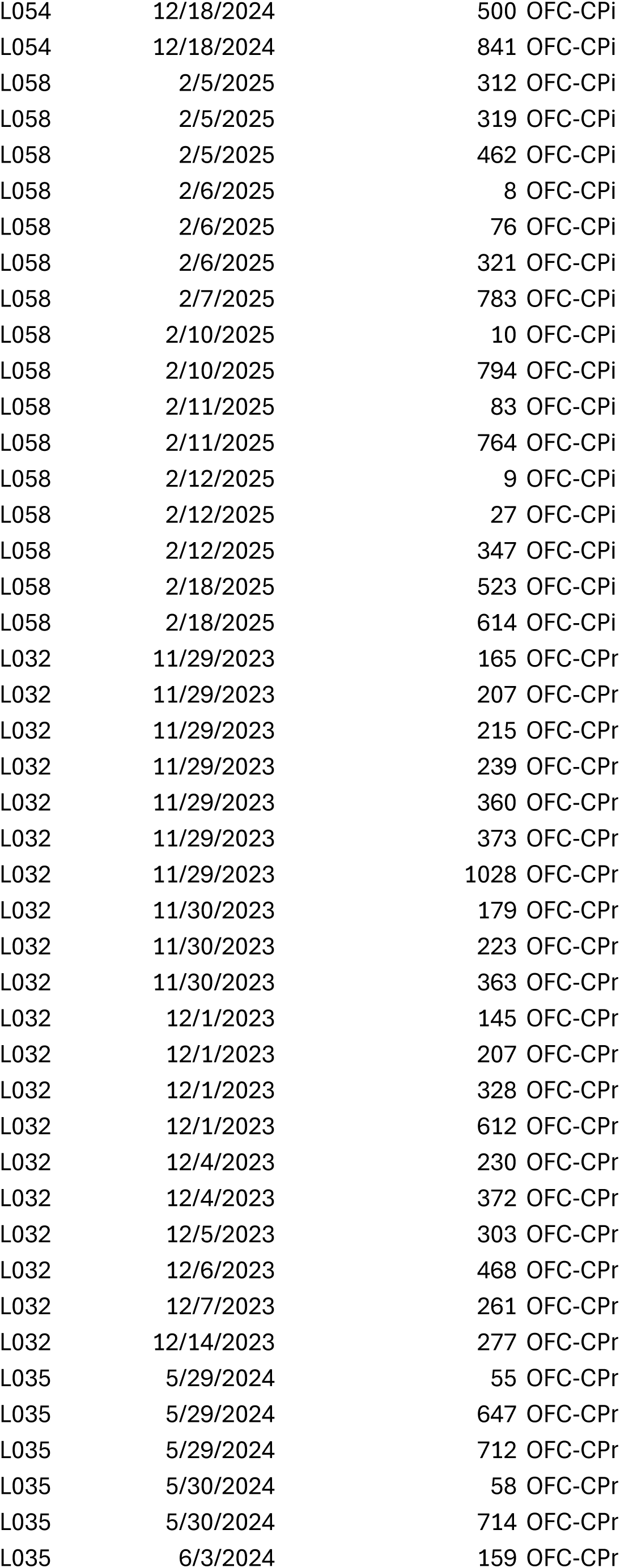
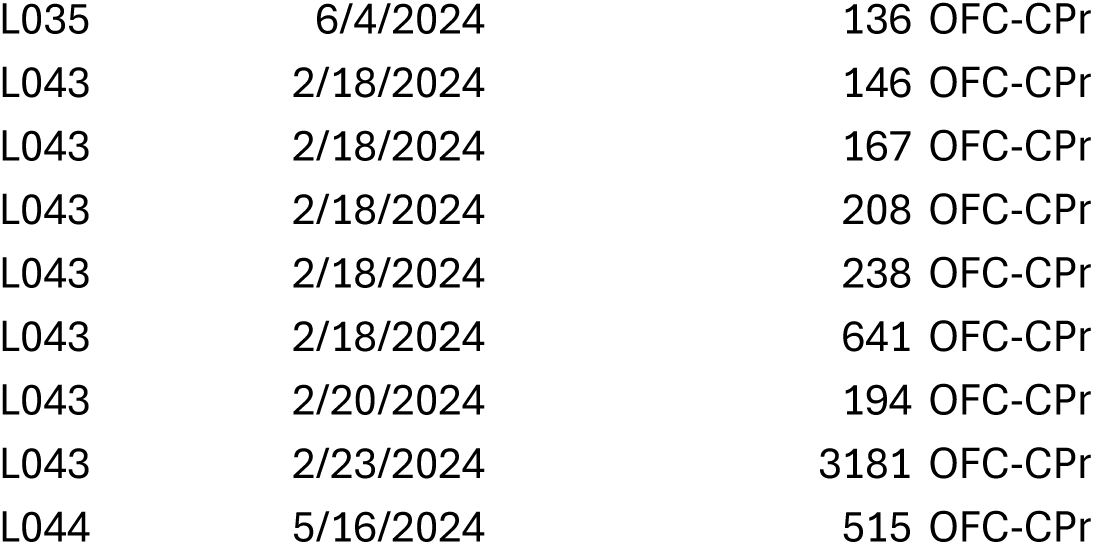

| **Rat ID** | **Total number of tagged cell** |
| --- | --- |
| 'L023' | 3 |
| 'L031' | 21 |
| 'L040' | 5 |
| 'L054' | 15 |
| 'L058' | 16 |
| L032 | 20 |
| L035 | 7 |
| L043 | 7 |
| L044 | 1 |

|  | OFC-CPi p-values |  |  |  | OCF-CPr p-values |  |  |  |
| --- | --- | --- | --- | --- | --- | --- | --- | --- |
| Time point | Offer cue | Reward Port on | Reward announced | Reward delivered | Offer cue | Reward Port on | Reward announced | Reward delivered |
| -1 | 0.34 | 0.199 | <0.001 | 0.015 | <0.001 | <0.001 | <0.001 | 0.052 |
| -0.9 | 0.006 | 0.165 | 0.043 | 0.289 | <0.001 | <0.001 | <0.001 | 0.003 |
| <0.001 | <0.001 | <0.001 | 0.194 | 0.034 | 0.003 | <0.001 | <0.001 | <0.001 |
| -0.7 | <0.001 | <0.001 | 0.002 | 0.002 | <0.001 | <0.001 | <0.001 | 0.16 |
| -0.6 | 0.142 | <0.001 | 0.077 | 0.43 | <0.001 | <0.001 | <0.001 | 0.01 |
| -0.5 | 0.119 | <0.001 | 0.001 | <0.001 | 0.007 | <0.001 | <0.001 | <0.001 |
| -0.4 | 0.007 | 0.001 | <0.001 | <0.001 | <0.001 | 0.237 | 0.006 | <0.001 |
| -0.3 | 0.002 | 0.032 | 0.05 | <0.001 | <0.001 | <0.001 | <0.001 | <0.001 |
| -0.2 | <0.001 | 0.415 | 0.301 | <0.001 | 0.007 | <0.001 | 0.036 | <0.001 |
| -0.1 | <0.001 | 0.091 | 0.003 | <0.001 | <0.001 | <0.001 | <0.001 | <0.001 |
| 0 | <0.001 | 0.254 | <0.001 | 0.367 | 0.053 | <0.001 | <0.001 | <0.001 |
| 0.1 | <0.001 | <0.001 | <0.001 | 0.42 | 0.003 | <0.001 | <0.001 | 0.046 |
| 0.2 | 0.472 | 0.097 | <0.001 | <0.001 | <0.001 | <0.001 | <0.001 | <0.001 |
| 0.3 | 0.374 | 0.168 | <0.001 | <0.001 | <0.001 | <0.001 | <0.001 | <0.001 |
| 0.4 | 0.013 | 0.097 | <0.001 | <0.001 | <0.001 | <0.001 | <0.001 | <0.001 |
| 0.5 | <0.001 | 0.252 | <0.001 | <0.001 | <0.001 | <0.001 | <0.001 | <0.001 |
| 0.6 | <0.001 | 0.166 | <0.001 | <0.001 | <0.001 | <0.001 | 0.038 | <0.001 |
| 0.7 | 0.059 | <0.001 | <0.001 | <0.001 | <0.001 | <0.001 | 0.04 | <0.001 |
| 0.8 | 0.03 | <0.001 | <0.001 | <0.001 | <0.001 | <0.001 | 0.002 | <0.001 |
| 0.9 | 0.084 | 0.022 | <0.001 | <0.001 | <0.001 | 0.262 | 0.003 | <0.001 |
| 1 | 0.001 | <0.001 | <0.001 | <0.001 | <0.001 | 0.057 | <0.001 | <0.001 |
| 1.1 | 0.027 | <0.001 | <0.001 | 0.003 | <0.001 | 0.137 | <0.001 | <0.001 |
| 1.2 | 0.011 | <0.001 | 0.035 | 0.51 | <0.001 | <0.001 | <0.001 | <0.001 |
| 1.3 | 0.31 | 0.006 | 0.203 | <0.001 | <0.001 | <0.001 | <0.001 | <0.001 |
| 1.4 | 0.268 | 0.056 | 0.092 | <0.001 | 0.001 | 0.002 | <0.001 | <0.001 |
| 1.5 | <0.001 | 0.098 | 0.044 | <0.001 | <0.001 | 0.001 | <0.001 | <0.001 |
| 1.6 | <0.001 | 0.08 | 0.161 | 0.002 | 0.003 | <0.001 | 0.026 | <0.001 |
| 1.7 | <0.001 | 0.254 | 0.001 | <0.001 | <0.001 | <0.001 | <0.001 | <0.001 |
| 1.8 | 0.003 | 0.315 | <0.001 | <0.001 | <0.001 | <0.001 | 0.002 | <0.001 |
| 1.9 | 0.23 | 0.114 | <0.001 | <0.001 | <0.001 | <0.001 | <0.001 | <0.001 |
| 2 | <0.001 | 0.466 | <0.001 | <0.001 | <0.001 | <0.001 | <0.001 | <0.001 |
| 2.1 | 0.006 | 0.078 | 0.03 | <0.001 | 0.008 | <0.001 | <0.001 | <0.001 |
| 2.2 | 0.08 | <0.001 | <0.001 | <0.001 | <0.001 | 0.324 | <0.001 | <0.001 |
| 2.3 | 0.026 | <0.001 | <0.001 | <0.001 | <0.001 | <0.001 | <0.001 | <0.001 |
| 2.4 | 0.076 | <0.001 | <0.001 | <0.001 | <0.001 | 0.039 | <0.001 | <0.001 |
| 2.5 | 0.143 | 0.147 | <0.001 | 0.026 | 0.002 | <0.001 | <0.001 | <0.001 |
| 2.6 | 0.001 | 0.021 | 0.018 | <0.001 | 0.173 | <0.001 | <0.001 | <0.001 |
| 2.7 | <0.001 | 0.39 | 0.001 | <0.001 | <0.001 | <0.001 | <0.001 | <0.001 |
| 2.8 | <0.001 | 0.151 | 0.005 | <0.001 | 0.085 | <0.001 | <0.001 | 0.365 |
| 2.9 | <0.001 | 0.049 | <0.001 | 0.006 | <0.001 | <0.001 | 0.001 | 0.034 |
| 3 | 0.039 | 0.068 | <0.001 | 0.001 | 0.5 | 0.294 | 0.051 | 0.457 |

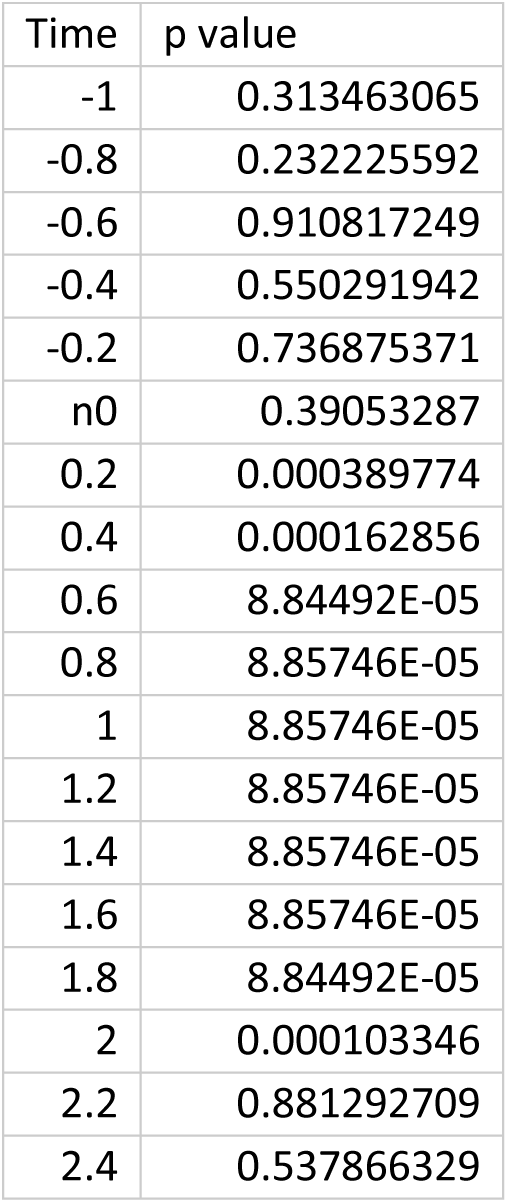

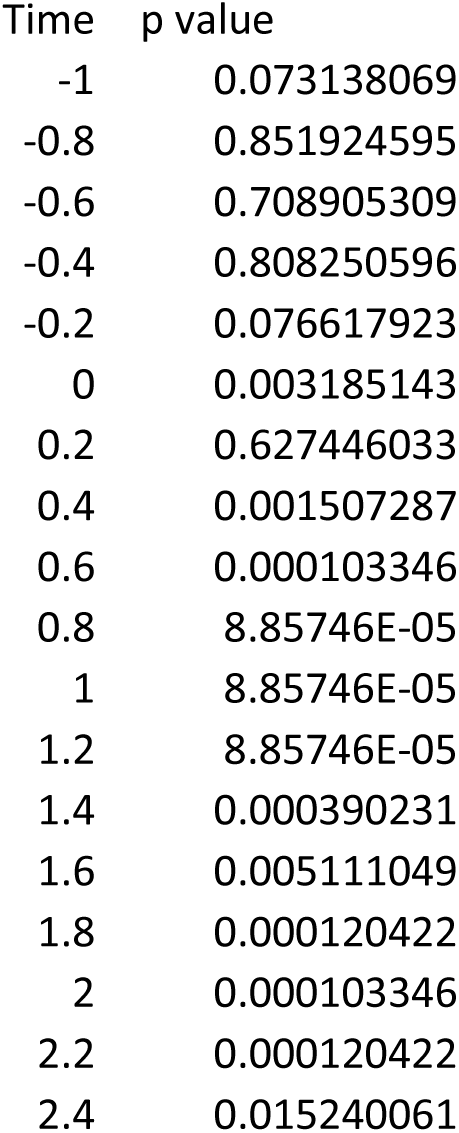

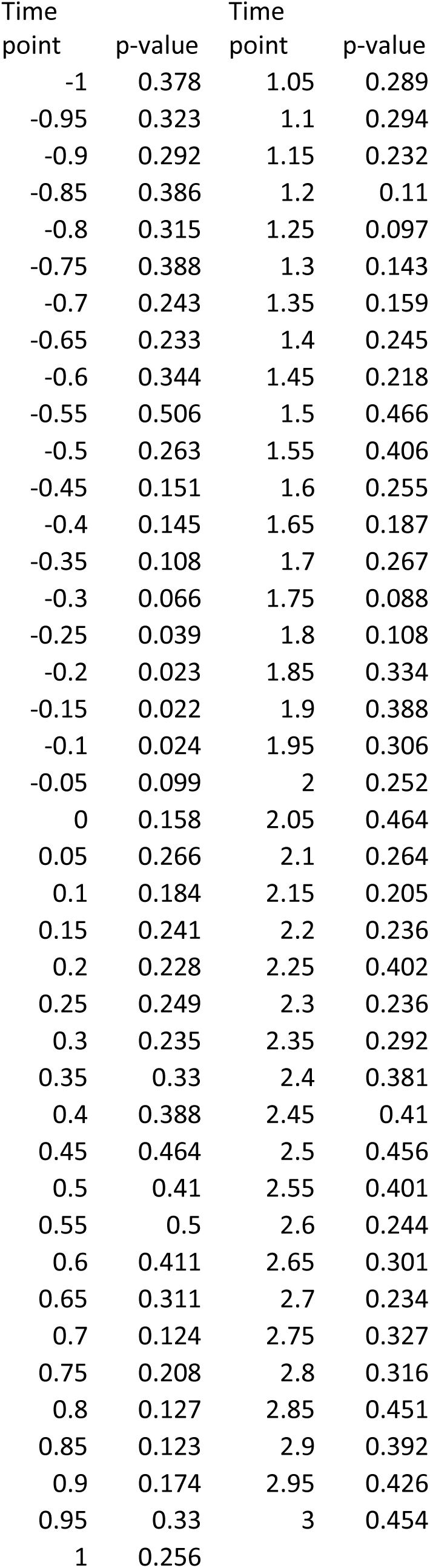

| Time | p value | Time | p value |
| --- | --- | --- | --- |
| -1 | 0.078 | 1.05 | 0 |
| -0.95 | 0.082 | 1.1 | 0 |
| -0.9 | 0.032 | 1.15 | 0 |
| -0.85 | 0.014 | 1.2 | 0 |
| -0.8 | 0.005 | 1.25 | 0.002 |
| -0.75 | 0.002 | 1.3 | 0.008 |
| -0.7 | 0.008 | 1.35 | 0.001 |
| -0.65 | 0.002 | 1.4 | 0.004 |
| -0.6 | 0.012 | 1.45 | 0.006 |
| -0.55 | 0.014 | 1.5 | 0.009 |
| -0.5 | 0.011 | 1.55 | 0.014 |
| -0.45 | 0.014 | 1.6 | 0.006 |
| -0.4 | 0.012 | 1.65 | 0.004 |
| -0.35 | 0.006 | 1.7 | 0.011 |
| -0.3 | 0.007 | 1.75 | 0.016 |
| -0.25 | 0.015 | 1.8 | 0.011 |
| -0.2 | 0.025 | 1.85 | 0.015 |
| -0.15 | 0.075 | 1.9 | 0.006 |
| -0.1 | 0.094 | 1.95 | 0.001 |
| -0.05 | 0.116 | 2 | 0.002 |
| 0 | 0.044 | 2.05 | 0.01 |
| 0.05 | 0.018 | 2.1 | 0.03 |
| 0.1 | 0.013 | 2.15 | 0.034 |
| 0.15 | 0 | 2.2 | 0.037 |
| 0.2 | 0.002 | 2.25 | 0.02 |
| 0.25 | 0.001 | 2.3 | 0.02 |
| 0.3 | 0 | 2.35 | 0.012 |
| 0.35 | 0 | 2.4 | 0.008 |
| 0.4 | 0.003 | 2.45 | 0.001 |
| 0.45 | 0.005 | 2.5 | 0 |
| 0.5 | 0.007 | 2.55 | 0.001 |
| 0.55 | 0.006 | 2.6 | 0.002 |
| 0.6 | 0.01 | 2.65 | 0 |
| 0.65 | 0.008 | 2.7 | 0 |
| 0.7 | 0.018 | 2.75 | 0 |
| 0.75 | 0.018 | 2.8 | 0.001 |
| 0.8 | 0.011 | 2.85 | 0.003 |
| 0.85 | 0.003 | 2.9 | 0.002 |
| 0.9 | 0.006 | 2.95 | 0.003 |
| 0.95 | 0 | 3 | 0 |
| 1 | 0 |  |  |
| -1 | 0.322 | 1.05 | 0.462 |
| -0.95 | 0.424 | 1.1 | 0.473 |
| -0.9 | 0.488 | 1.15 | 0.444 |
| -0.85 | 0.26 | 1.2 | 0.428 |
| -0.8 | 0.171 | 1.25 | 0.44 |
| -0.75 | 0.102 | 1.3 | 0.485 |
| -0.7 | 0.041 | 1.35 | 0.412 |
| -0.65 | 0.042 | 1.4 | 0.454 |
| -0.6 | 0.119 | 1.45 | 0.453 |
| -0.55 | 0.13 | 1.5 | 0.467 |
| -0.5 | 0.097 | 1.55 | 0.442 |
| -0.45 | 0.062 | 1.6 | 0.411 |
| -0.4 | 0.085 | 1.65 | 0.355 |
| -0.35 | 0.077 | 1.7 | 0.252 |
| -0.3 | 0.159 | 1.75 | 0.225 |
| -0.25 | 0.263 | 1.8 | 0.271 |
| -0.2 | 0.343 | 1.85 | 0.338 |
| -0.15 | 0.44 | 1.9 | 0.412 |
| -0.1 | 0.437 | 1.95 | 0.421 |
| -0.05 | 0.343 | 2 | 0.315 |
| 0 | 0.493 | 2.05 | 0.34 |
| 0.05 | 0.192 | 2.1 | 0.383 |
| 0.1 | 0.181 | 2.15 | 0.323 |
| 0.15 | 0.187 | 2.2 | 0.406 |
| 0.2 | 0.105 | 2.25 | 0.417 |
| 0.25 | 0.081 | 2.3 | 0.475 |
| 0.3 | 0.108 | 2.35 | 0.453 |
| 0.35 | 0.227 | 2.4 | 0.417 |
| 0.4 | 0.255 | 2.45 | 0.396 |
| 0.45 | 0.305 | 2.5 | 0.389 |
| 0.5 | 0.271 | 2.55 | 0.439 |
| 0.55 | 0.38 | 2.6 | 0.33 |
| 0.6 | 0.416 | 2.65 | 0.318 |
| 0.65 | 0.417 | 2.7 | 0.339 |
| 0.7 | 0.437 | 2.75 | 0.341 |
| 0.75 | 0.497 | 2.8 | 0.312 |
| 0.8 | 0.393 | 2.85 | 0.402 |
| 0.85 | 0.396 | 2.9 | 0.392 |
| 0.9 | 0.415 | 2.95 | 0.308 |
| 0.95 | 0.417 | 3 | 0.26 |
| 1 | 0.403 |  |  |
| -1 | 0.357 | 1.05 | 0.02 |
| -0.95 | 0.469 | 1.1 | 0.013 |
| -0.9 | 0.449 | 1.15 | 0.019 |
| -0.85 | 0.413 | 1.2 | 0.006 |
| -0.8 | 0.489 | 1.25 | 0.003 |
| -0.75 | 0.389 | 1.3 | 0 |
| -0.7 | 0.427 | 1.35 | 0.002 |
| -0.65 | 0.436 | 1.4 | 0.005 |
| -0.6 | 0.406 | 1.45 | 0 |
| -0.55 | 0.253 | 1.5 | 0.002 |
| -0.5 | 0.153 | 1.55 | 0.004 |
| -0.45 | 0.278 | 1.6 | 0.011 |
| -0.4 | 0.285 | 1.65 | 0.007 |
| -0.35 | 0.408 | 1.7 | 0.015 |
| -0.3 | 0.471 | 1.75 | 0.006 |
| -0.25 | 0.486 | 1.8 | 0.001 |
| -0.2 | 0.389 | 1.85 | 0 |
| -0.15 | 0.299 | 1.9 | 0.002 |
| -0.1 | 0.187 | 1.95 | 0.001 |
| -0.05 | 0.117 | 2 | 0 |
| 0 | 0.09 | 2.05 | 0 |
| 0.05 | 0.047 | 2.1 | 0 |
| 0.1 | 0.051 | 2.15 | 0 |
| 0.15 | 0.017 | 2.2 | 0 |
| 0.2 | 0.011 | 2.25 | 0 |
| 0.25 | 0.013 | 2.3 | 0 |
| 0.3 | 0.014 | 2.35 | 0 |
| 0.35 | 0.007 | 2.4 | 0 |
| 0.4 | 0.011 | 2.45 | 0.001 |
| 0.45 | 0.007 | 2.5 | 0 |
| 0.5 | 0.009 | 2.55 | 0 |
| 0.55 | 0.012 | 2.6 | 0.001 |
| 0.6 | 0.012 | 2.65 | 0.002 |
| 0.65 | 0.017 | 2.7 | 0.002 |
| 0.7 | 0.019 | 2.75 | 0.005 |
| 0.75 | 0.006 | 2.8 | 0.006 |
| 0.8 | 0.009 | 2.85 | 0.004 |
| 0.85 | 0.009 | 2.9 | 0.009 |
| 0.9 | 0.011 | 2.95 | 0.006 |
| 0.95 | 0.006 | 3 | 0 |
| 1 | 0.032 |  |  |
| -1 | 0.284 | 1.05 | 0.392 |
| -0.95 | 0.362 | 1.1 | 0.5 |
| -0.9 | 0.258 | 1.15 | 0.439 |
| -0.85 | 0.164 | 1.2 | 0.292 |
| -0.8 | 0.25 | 1.25 | 0.301 |
| -0.75 | 0.392 | 1.3 | 0.288 |
| -0.7 | 0.379 | 1.35 | 0.194 |
| -0.65 | 0.418 | 1.4 | 0.155 |
| -0.6 | 0.414 | 1.45 | 0.15 |
| -0.55 | 0.173 | 1.5 | 0.143 |
| -0.5 | 0.071 | 1.55 | 0.117 |
| -0.45 | 0.031 | 1.6 | 0.123 |
| -0.4 | 0.071 | 1.65 | 0.15 |
| -0.35 | 0.179 | 1.7 | 0.105 |
| -0.3 | 0.342 | 1.75 | 0.045 |
| -0.25 | 0.477 | 1.8 | 0.052 |
| -0.2 | 0.339 | 1.85 | 0.05 |
| -0.15 | 0.275 | 1.9 | 0.069 |
| -0.1 | 0.364 | 1.95 | 0.101 |
| -0.05 | 0.412 | 2 | 0.113 |
| 0 | 0.403 | 2.05 | 0.116 |
| 0.05 | 0.077 | 2.1 | 0.139 |
| 0.1 | 0.01 | 2.15 | 0.066 |
| 0.15 | 0.017 | 2.2 | 0.04 |
| 0.2 | 0.074 | 2.25 | 0.034 |
| 0.25 | 0.103 | 2.3 | 0.021 |
| 0.3 | 0.15 | 2.35 | 0.012 |
| 0.35 | 0.325 | 2.4 | 0.011 |
| 0.4 | 0.427 | 2.45 | 0.016 |
| 0.45 | 0.452 | 2.5 | 0.018 |
| 0.5 | 0.43 | 2.55 | 0.01 |
| 0.55 | 0.469 | 2.6 | 0.003 |
| 0.6 | 0.475 | 2.65 | 0.004 |
| 0.65 | 0.414 | 2.7 | 0.009 |
| 0.7 | 0.422 | 2.75 | 0.005 |
| 0.75 | 0.47 | 2.8 | 0.013 |
| 0.8 | 0.429 | 2.85 | 0.018 |
| 0.85 | 0.359 | 2.9 | 0.035 |
| 0.9 | 0.318 | 2.95 | 0.047 |
| 0.95 | 0.431 | 3 | 0.026 |
| 1 | 0.38 |  |  |
| -1 | 0.186 | 1.05 | <0.001 |
| -0.95 | 0.084 | 1.1 | <0.001 |
| -0.9 | 0.044 | 1.15 | <0.001 |
| -0.85 | 0.022 | 1.2 | 0.001 |
| -0.8 | 0.004 | 1.25 | 0.001 |
| -0.75 | 0.005 | 1.3 | 0.004 |
| -0.7 | 0.004 | 1.35 | <0.001 |
| -0.65 | 0.001 | 1.4 | 0.004 |
| -0.6 | 0.012 | 1.45 | 0.005 |
| -0.55 | 0.017 | 1.5 | 0.009 |
| -0.5 | 0.015 | 1.55 | 0.021 |
| -0.45 | 0.022 | 1.6 | 0.023 |
| -0.4 | 0.008 | 1.65 | 0.007 |
| -0.35 | 0.003 | 1.7 | 0.012 |
| -0.3 | 0.002 | 1.75 | 0.009 |
| -0.25 | 0.018 | 1.8 | 0.003 |
| -0.2 | 0.014 | 1.85 | 0.01 |
| -0.15 | 0.065 | 1.9 | 0.005 |
| -0.1 | 0.065 | 1.95 | 0.002 |
| -0.05 | 0.092 | 2 | 0.002 |
| 0 | 0.029 | 2.05 | 0.018 |
| 0.05 | 0.013 | 2.1 | 0.029 |
| 0.1 | 0.009 | 2.15 | 0.026 |
| 0.15 | 0.002 | 2.2 | 0.027 |
| 0.2 | <0.001 | 2.25 | 0.018 |
| 0.25 | <0.001 | 2.3 | 0.016 |
| 0.3 | 0.003 | 2.35 | 0.01 |
| 0.35 | 0.002 | 2.4 | 0.004 |
| 0.4 | 0.003 | 2.45 | 0.002 |
| 0.45 | <0.001 | 2.5 | 0.002 |
| 0.5 | 0.001 | 2.55 | 0.001 |
| 0.55 | 0.003 | 2.6 | 0.001 |
| 0.6 | 0.015 | 2.65 | <0.001 |
| 0.65 | 0.004 | 2.7 | <0.001 |
| 0.7 | 0.006 | 2.75 | 0.001 |
| 0.75 | 0.008 | 2.8 | 0.002 |
| 0.8 | 0.004 | 2.85 | 0.003 |
| 0.85 | 0.004 | 2.9 | <0.001 |
| 0.9 | 0.004 | 2.95 | 0.001 |
| 0.95 | <0.001 | 3 | 0.001 |
| 1 | <0.001 |  |  |
| -1 | 0.39 | 1.05 | 0.463 |
| -0.95 | 0.486 | 1.1 | 0.466 |
| -0.9 | 0.491 | 1.15 | 0.477 |
| -0.85 | 0.219 | 1.2 | 0.496 |
| -0.8 | 0.139 | 1.25 | 0.49 |
| -0.75 | 0.105 | 1.3 | 0.501 |
| -0.7 | 0.046 | 1.35 | 0.46 |
| -0.65 | 0.024 | 1.4 | 0.486 |
| -0.6 | 0.108 | 1.45 | 0.463 |
| -0.55 | 0.141 | 1.5 | 0.482 |
| -0.5 | 0.098 | 1.55 | 0.461 |
| -0.45 | 0.072 | 1.6 | 0.425 |
| -0.4 | 0.089 | 1.65 | 0.372 |
| -0.35 | 0.087 | 1.7 | 0.261 |
| -0.3 | 0.204 | 1.75 | 0.204 |
| -0.25 | 0.312 | 1.8 | 0.284 |
| -0.2 | 0.321 | 1.85 | 0.326 |
| -0.15 | 0.361 | 1.9 | 0.447 |
| -0.1 | 0.401 | 1.95 | 0.351 |
| -0.05 | 0.238 | 2 | 0.303 |
| 0 | 0.321 | 2.05 | 0.305 |
| 0.05 | 0.22 | 2.1 | 0.31 |
| 0.1 | 0.191 | 2.15 | 0.322 |
| 0.15 | 0.206 | 2.2 | 0.359 |
| 0.2 | 0.095 | 2.25 | 0.385 |
| 0.25 | 0.089 | 2.3 | 0.44 |
| 0.3 | 0.124 | 2.35 | 0.444 |
| 0.35 | 0.225 | 2.4 | 0.456 |
| 0.4 | 0.206 | 2.45 | 0.396 |
| 0.45 | 0.249 | 2.5 | 0.39 |
| 0.5 | 0.262 | 2.55 | 0.39 |
| 0.55 | 0.365 | 2.6 | 0.368 |
| 0.6 | 0.396 | 2.65 | 0.332 |
| 0.65 | 0.428 | 2.7 | 0.303 |
| 0.7 | 0.441 | 2.75 | 0.331 |
| 0.75 | 0.484 | 2.8 | 0.31 |
| 0.8 | 0.411 | 2.85 | 0.387 |
| 0.85 | 0.403 | 2.9 | 0.375 |
| 0.9 | 0.39 | 2.95 | 0.3 |
| 0.95 | 0.451 | 3 | 0.218 |
| 1 | 0.414 |  |  |
| -1 | 0.344 | 1.05 | 0.039 |
| -0.95 | 0.377 | 1.1 | 0.052 |
| -0.9 | 0.327 | 1.15 | 0.015 |
| -0.85 | 0.314 | 1.2 | 0.016 |
| -0.8 | 0.319 | 1.25 | 0.006 |
| -0.75 | 0.358 | 1.3 | <0.001 |
| -0.7 | 0.223 | 1.35 | <0.001 |
| -0.65 | 0.201 | 1.4 | 0.003 |
| -0.6 | 0.222 | 1.45 | 0.003 |
| -0.55 | 0.166 | 1.5 | 0.002 |
| -0.5 | 0.102 | 1.55 | 0.005 |
| -0.45 | 0.123 | 1.6 | 0.006 |
| -0.4 | 0.192 | 1.65 | 0.004 |
| -0.35 | 0.224 | 1.7 | 0.001 |
| -0.3 | 0.202 | 1.75 | 0.002 |
| -0.25 | 0.333 | 1.8 | <0.001 |
| -0.2 | 0.299 | 1.85 | <0.001 |
| -0.15 | 0.227 | 1.9 | <0.001 |
| -0.1 | 0.108 | 1.95 | <0.001 |
| -0.05 | 0.049 | 2 | 0.001 |
| 0 | 0.031 | 2.05 | <0.001 |
| 0.05 | 0.014 | 2.1 | <0.001 |
| 0.1 | 0.003 | 2.15 | <0.001 |
| 0.15 | 0.002 | 2.2 | <0.001 |
| 0.2 | 0.003 | 2.25 | <0.001 |
| 0.25 | 0.007 | 2.3 | <0.001 |
| 0.3 | 0.002 | 2.35 | <0.001 |
| 0.35 | 0.002 | 2.4 | <0.001 |
| 0.4 | 0.001 | 2.45 | <0.001 |
| 0.45 | 0.004 | 2.5 | <0.001 |
| 0.5 | 0.005 | 2.55 | <0.001 |
| 0.55 | 0.01 | 2.6 | <0.001 |
| 0.6 | 0.018 | 2.65 | <0.001 |
| 0.65 | 0.03 | 2.7 | <0.001 |
| 0.7 | 0.031 | 2.75 | <0.001 |
| 0.75 | 0.021 | 2.8 | <0.001 |
| 0.8 | 0.021 | 2.85 | <0.001 |
| 0.85 | 0.023 | 2.9 | <0.001 |
| 0.9 | 0.033 | 2.95 | 0.004 |
| 0.95 | 0.036 | 3 | <0.001 |
| 1 | 0.095 |  |  |
| -1 | 0.446 | 1.05 | 0.15 |
| -0.95 | 0.118 | 1.1 | 0.167 |
| -0.9 | 0.055 | 1.15 | 0.096 |
| -0.85 | 0.034 | 1.2 | 0.077 |
| -0.8 | 0.087 | 1.25 | 0.12 |
| -0.75 | 0.107 | 1.3 | 0.14 |
| -0.7 | 0.094 | 1.35 | 0.172 |
| -0.65 | 0.203 | 1.4 | 0.139 |
| -0.6 | 0.231 | 1.45 | 0.112 |
| -0.55 | 0.063 | 1.5 | 0.112 |
| -0.5 | 0.144 | 1.55 | 0.095 |
| -0.45 | 0.047 | 1.6 | 0.072 |
| -0.4 | 0.065 | 1.65 | 0.07 |
| -0.35 | 0.345 | 1.7 | 0.057 |
| -0.3 | 0.457 | 1.75 | 0.033 |
| -0.25 | 0.45 | 1.8 | 0.048 |
| -0.2 | 0.312 | 1.85 | 0.046 |
| -0.15 | 0.298 | 1.9 | 0.076 |
| -0.1 | 0.401 | 1.95 | 0.061 |
| -0.05 | 0.442 | 2 | 0.077 |
| 0 | 0.436 | 2.05 | 0.072 |
| 0.05 | 0.184 | 2.1 | 0.047 |
| 0.1 | 0.063 | 2.15 | 0.03 |
| 0.15 | 0.018 | 2.2 | 0.022 |
| 0.2 | 0.026 | 2.25 | 0.027 |
| 0.25 | 0.037 | 2.3 | 0.035 |
| 0.3 | 0.058 | 2.35 | 0.04 |
| 0.35 | 0.103 | 2.4 | 0.036 |
| 0.4 | 0.28 | 2.45 | 0.027 |
| 0.45 | 0.26 | 2.5 | 0.029 |
| 0.5 | 0.203 | 2.55 | 0.016 |
| 0.55 | 0.225 | 2.6 | 0.007 |
| 0.6 | 0.235 | 2.65 | 0.009 |
| 0.65 | 0.194 | 2.7 | 0.016 |
| 0.7 | 0.248 | 2.75 | 0.025 |
| 0.75 | 0.329 | 2.8 | 0.009 |
| 0.8 | 0.364 | 2.85 | 0.017 |
| 0.85 | 0.308 | 2.9 | 0.045 |
| 0.9 | 0.241 | 2.95 | 0.055 |
| 0.95 | 0.284 | 3 | 0.053 |
| 1 | 0.224 |  |  |

